# Gwas and Genomic Selection For Increased Anthocyanin Content in Purple Corn

**DOI:** 10.1101/2020.05.20.107359

**Authors:** Laura A. Chatham, John A. Juvik

**Affiliations:** Department of Crop Sciences, University of Illinois at Urbana Champaign

**Keywords:** Anthocyanins, flavonoid biosynthesis pathway, flux, maize pericarp, purple corn, genotyping-by-sequencing, genomic selection

## Abstract

Purple corn offers an attractive source of economical natural anthocyanin-based colorant for use in foods and beverages. Yet to maximize the scalability and meet growing demands, both anthocyanin concentrations and agronomic performance must improve in purple corn varieties. We studied flux through the flavonoid biosynthesis pathway using GWAS data derived from a diverse purple corn landrace with anthocyanin-rich pericarp, Apache Red. Trends between flavonoid endpoints suggest that regulators of total flux into the pathway and regulators of partitioning within the pathway may both represent targets for maximizing anthocyanin content. A peak at the end of chromosome 1 near *Aat1 (Anthocyanin acyltransferase1*) was highly significant in all approaches taken to map anthocyanin flux, suggesting the structural modification of malonylation is required for maximal anthocyanin production. We also identified several candidate MATEs and H^+^ ATPases that could assist in the preferential transport of acylated anthocyanins into the vacuole. These and other candidates identified here suggest there is still much to learn about the regulation of anthocyanin biosynthesis in the pericarp of purple corn. The efficacy of genomic predictions in the population was also studied, yielding an accuracy of 0.71 with cross validation for total anthocyanin content with no improvement found when known anthocyanin regulators were added to the model. These data suggest that genomic selection could be employed effectively in a purple corn breeding program, and especially for a landrace improvement program.

## Introduction

With continued trends in consumer preference for less processed foods, rising demand for naturally pigmented food and beverages necessitates the development of economical natural colorant sources. Purple corn is a good source of anthocyanins, the water soluble, red to purple pigments found in many types of fruits and vegetables. As a scalable commodity, corn could provide a natural colorant source capable of meeting growing consumer demands. However, widescale production will require breeding to create lines with maximum anthocyanin production while maintaining yield and other agronomic traits.

Various strategies can be taken in breeding for increasing the economic feasibility of using purple corn to replace artificial colorants. Ultimately the goal is to increase anthocyanin yield per unit input and ideally both per-kernel anthocyanin content and per-acre yield could be increased to compound the total anthocyanin yield per acre. Accomplishing this will require identifying the highest anthocyanin producing purple corn lines and backcrossing into current elite inbreds. With each round of backcrossing, selections must be made to minimize linkage drag while avoiding the loss of any factors associated with maximal anthocyanin production. Anthocyanin biosynthesis involves a number of different structural genes and regulatory mechanisms, all of which have the potential to become unfixed alleles upon backcrossing with non-pigmented lines. Loss of any of these factors could influence anthocyanin yield to varying degrees.

Nearly all of the anthocyanin structural genes have been identified with the exception of those associated with anthocyanin decorations post-synthesis^1^. While the major regulatory players have been identified, current models in maize are simplistic compared to those in other anthocyanin producing species^2^. The anthocyanin MBW regulatory complex consists of an R2R3-MYB protein, a bHLH (basic helix-loop-helix) protein, and WDR (WD-repeat) domain protein^3^. Typically, *Colored aleurone1* (*C1*, MYB), *Colored1* (*R1*, bHLH), and *Pale aleurone color1* (*Pac1*, WDR) together create the regulatory complex and activate structural genes in the kernel aleurone. A bHLH-like recessive intensifier *in1* (*intensifier1*) with similarity to *R1*, increases anthocyanin pigmentation in aleurone^4^. More recently an EMSY-like factor, *RIF1* (*R interacting factor1*) was found to act as a switch to help control activation of structural genes^5^. An analogous MBW complex likely forms to regulate anthocyanin biosynthesis in plant tissue and kernel pericarp. *Booster1*(*B1*) and *Plant color1* (*Pl1*) are the bHLH and MYB regulatory factors, respectively, most often associated with regulation in plant tissues. However, there are exceptions in which certain alleles of *R1* (e.g. *R1-ch)* operate in pericarp and certain *B1* alleles (e.g. *B1-Peru*) operate in aleurone^6^. Whether the WDR, *RIF1*-like, or *IN1*-like proteins function in pericarp has yet to be determined, and no homologous pericarp versions of these have been identified either. Furthermore, R2R3-MYB repressors of anthocyanin biosynthesis have been identified in a number of other species, yet none have been identified in maize^7^. MYB repressors associated with the phenylpropanoid pathway and downstream lignin biosynthesis have been identified^8^.

This regulation of the structural genes occurs in the nucleus and the structural gene products function together in a flavonoid metabolon at the cytoplasmic face of the endoplasmic reticulum^9^. Anthocyanins are then transported to the vacuole for storage. In maize, *Bz2* (*Bronze2*) encodes a glutathione S-transferase which assists in anthocyanin transport and is required to prevent anthocyanin oxidation, resulting in a bronze colored kernel^10^. *Mrp3* (*Multidrug resistance-associated protein3*) produces an ABC transporter that shuttles anthocyanins into the vacuole^11^. Both were identified and studied in aleurone-pigmented lines. In other anthocyanin-producing species two other mechanisms of vacuolar localization have been identified in addition to the GST/ABC transporter system – MATE-transporters and vesicle trafficking. MATE-transporters, either as H^+^/flavonoid antiporters or in combination with other vacuolar proton gradient maintaining mechanisms have been identified in *Arabidopsis, Medicago*, tomato *(S. lycopersicum*) and radish (*R. sativa*)^12–17^. Support for vesicle trafficking of anthocyanins and a subsequent autophagy-like mechanism of vacuolar transport has been found in *Arabidopsis, Brassica napus*, and grapevine (*V. vinifera)*. Some evidence for this mechanism exists in maize. Anthocyanin-containing vesicles and anthocyanin vacuolar inclusions (AVIs) were shown to fuse in response to light^18^

In addition to anthocyanins, several other flavonoids are present in maize. Phlobaphenes, orange to brick red pigments accumulating in cob and pericarp, are polymers of the 3-deoxyflavonoids, luteoforol or apiforol, and share the early steps of their biosynthesis and thus carbon flux with anthocyanins. The structural genes responsible for phlobaphene biosynthesis are controlled by *Pericarp color1* (*P1*), an R2R3 MYB similar to *C1*/*Pl1*. In contrast to the anthocyanin regulating MYBs (*C1/Pl1), P1* functions independently, not requiring a bHLH partner for regulatory activity. *P1* also regulates C-glycosyl flavone biosynthesis, a group of compounds that includes maysin, an insecticidal compound conferring resistance to corn earworm (*Helicoverpa zea*)^19^.

With multiple endpoints, branch points and shared precursors, flux through the flavonoid pathway in maize is complex. Breeding for maximal anthocyanin content is likely to benefit greatly from a better understanding of the mechanisms regulating this flux. Furthermore, the identification of additional factors associated with anthocyanin content may help prevent their loss during backcrossing and enable marker assisted selection to expedite the process. Here we explore flux through the flavonoid pathway in a landrace with variability in anthocyanin, flavone, and flavonol content and candidate genes underlying this variability. Given the complexity and quantitative nature of the trait, we also gauge the accuracy of genomic selection with and without major regulatory players.

## Methods

### Plant materials

Apache Red seeds (S_0_) were purchased from Siskiyou Seeds (Williams, OR) and lines were created as described and illustrated in this article’s companion article (Chatham and Juvik, submitted to G3). Briefly, each round of selection involved phenotyping (HPLC) and selecting lines with anthocyanin content only and with the most variability in anthocyanin composition. This produced 181 S_2_ lines that were used for a pilot round of genotyping and phenotyping. After further breeding and selection, 1148 S_2_, S_3_, and S_4_ lines were produced that were used for genotyping and phenotyping. Selected S_2_, S_3_, and S4 seeds were sewn the following season and bulked by plot to produce seed for use in other collaborative projects. Kernels from S_3_, S_4_, and S_5_ plots were phenotyped as described below.

### Extractions, HPLC analysis, and UV-visible spectroscopy

All anthocyanin/flavone phenotyping, including extraction, HPLC analysis, and UV-visible spectroscopy were performed as described previously (in this article’s companion). This reference also describes seedling germination, growth, and sampling for DNA extraction. Variables were transformed as needed for comparisons.

### Genotyping and SNP discovery

GBS libraries were prepared as described previously and sequencing was performed by the Roy J. Carver Biotechnology Center at the University of Illinois in Urbana-Champaign (Chatham and Juvik, submitted to G3). The 1148 lines (S_2_, S_3_, and S_4_ selfed progeny) were sequenced on three lanes of Illumina HiSeq 4000 with single-end 100 nucleotide reads and multiplexed to obtain approximately equal numbers of reads on each lane. SNP discovery was carried out using the TASSEL 5.0 GBS v2 pipeline^20^ as described previously (Chatham and Juvik, submitted to G3)

### Association mapping and genomic selection

The R GAPIT (Genome Association and Prediction Integrated Tool) package^21^ was used for association mapping, and the RRBlup package^22^ was used for genomic selection. GWAS was also run individually for each chromosome using a leave-one-out method of calculating kinship to retain maximal power for detecting significant associations^23^. For genomic selection, prediction accuracy averages were measured from 25 replicates of five-fold cross validation.

## Results and Discussion

### Apache red population

Phenotypes were generated at each stage of the Apache Red (AR) population development and used to calculate heritability from parent offspring correlations^24^. Anthocyanin content heritability ranged from 0.44 to 0.57, suggesting improvement can be made through selection. Figure 1 shows a summary of kernel colors obtained from the AR population. AR S_0_ seeds were all dark purple, indicative of anthocyanins in pericarp, but S_1_s varied in color. Most were dark purple, anthocyanin-containing, but we observed a range of water insoluble phlobaphene-containing kernels colored from orange to purple-red and some kernels that were colorless or had minimal anthocyanin or phlobaphene pigmentation. While selection for highly anthocyanin-pigmented kernels appears to have been successful given the decrease in the proportion of colorless or phlobaphene containing kernels across generations of self-pollination, the presence of colorless and phlobaphene containing kernels by the S_4_ (S_3_ pericarp) generation suggests that anthocyanin content is controlled by multiple segregating loci in this population. An alternative explanation is that some of the segregating alleles could be paramutable, spontaneously converting to paramutagenic alleles, which then heritably silence other paramutable alleles^25^. Even in the most advanced lines from the last generations grown (S_3_s, S_4_s, and S_5_s), which were intended to bulk seed for other collaborative projects on anthocyanin-rich maize, colorless and phlobaphene containing ears were still present and had to be discarded at the time of harvest.

**Figure 1:**
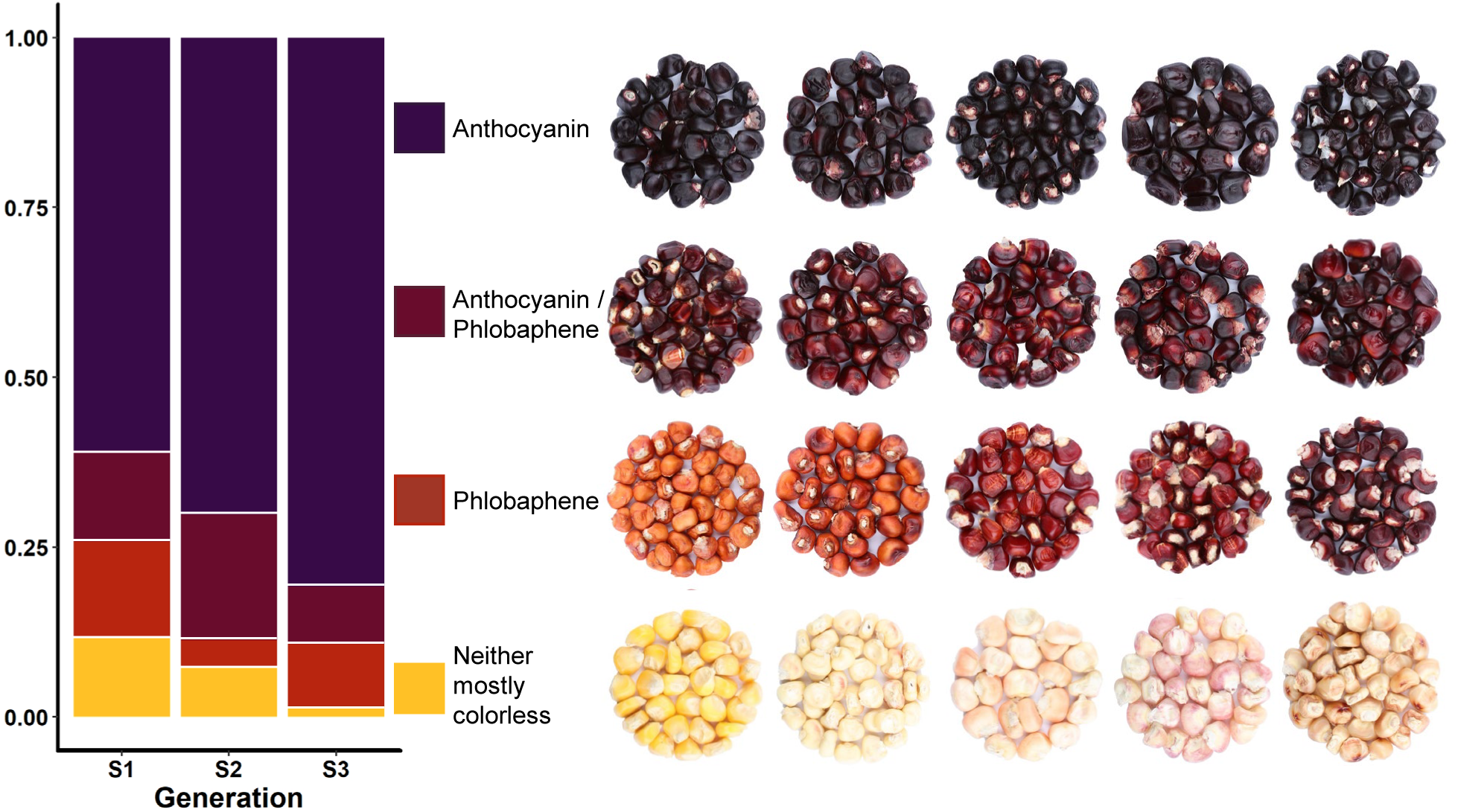
Proportions of lines from each generation containing anthocyanin and phlobaphenes based on visual assessment. Kernel pictures show ranges for each category. Generations are labeled based on maternal pericarp (i.e. S2, S3, and S4 plants have pericarp have maternal pericarp representative the S1, S2, and S3 generations, respectively)

### Phenotyping flux

To understand the factors influencing total anthocyanin content we compared the relationship between proportions of flavonoid end products and total anthocyanin content. Several simplistic models of flux can be considered. Under a finite flux, zero-sum hypothesis, one would expect anthocyanin content to decline as proportions of other flavonoids increase. Conversely if pathways are regulated independently or are regulated together by some universal regulator, one would expect to see no correlation or positive correlations between anthocyanin content and proportions of other flavonoids, respectively.

Flavone and anthocyanin content were positively correlated (r = 0.28, p < 2.2e-16) (Figure 2A), suggesting a major regulatory factor functioning at a step prior to the branch point between anthocyanins and flavones. Conversely, the proportion of anthocyanins present as flavonol-anthocyanin condensed forms was negatively correlated with total anthocyanin content (r = −0.27, p = 6.24e-16) (Figure 2B), lending support to a competitive zero-sum-like model of flux for these forms. Correlations with specific anthocyanin types were also examined. Proportion of acylated anthocyanins was positively correlated with anthocyanin content (r = 0.23, p = 1.51e-14) (Figure 2C), and the proportion of cyanidin-derived anthocyanins was negatively associated with anthocyanin content (r = −0.31, p < 2.2e-16). These correlations could be an indication of substrate specificity in downstream processes or could be due to linkage or population structure. All lines here were derived from the AR landrace, making population structure an important consideration.

**Figure 2:**
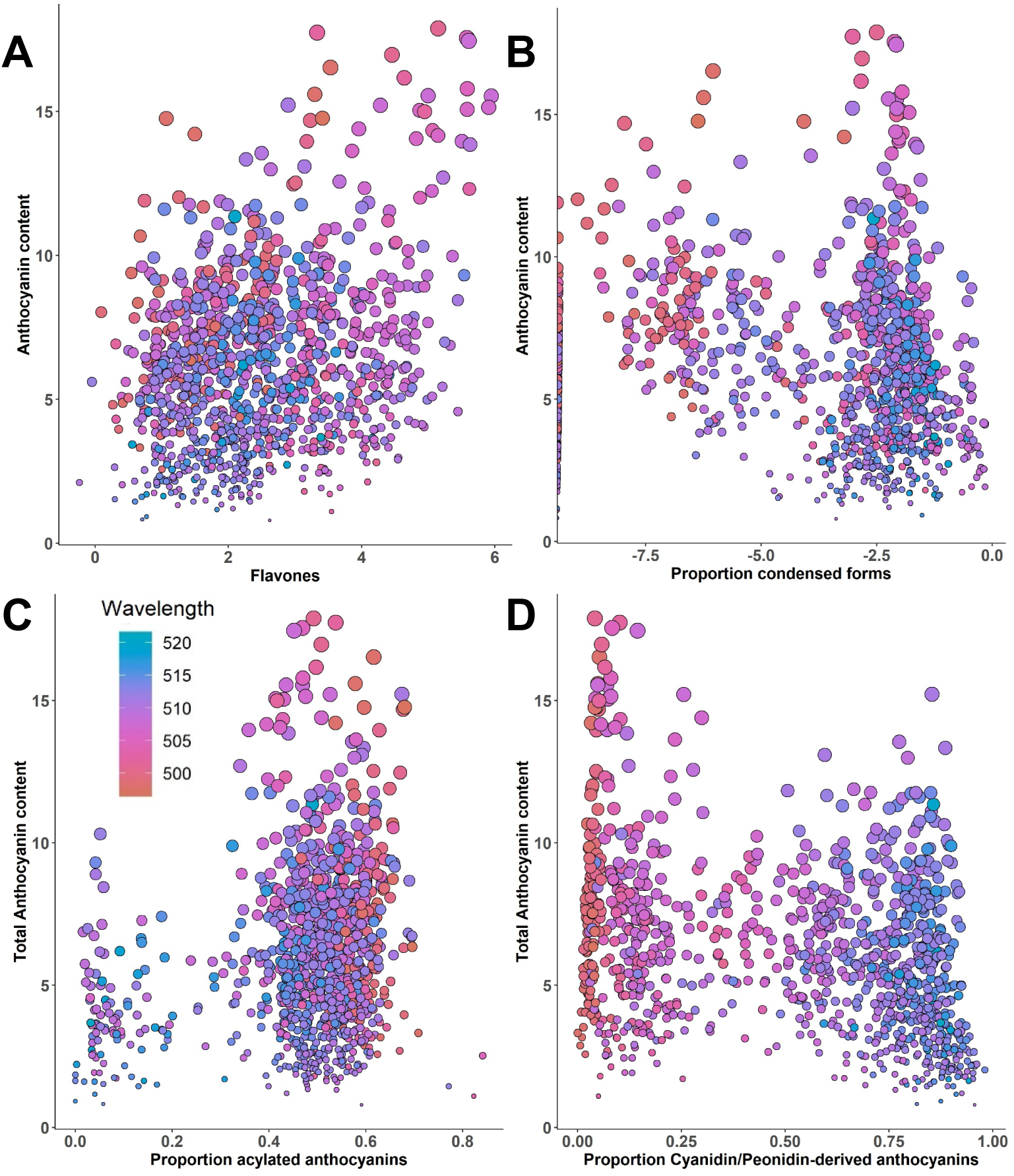
Effects of flavone content (A, log transformed, μg/ml), and proportions of condensed (B, log transformed), acylated (C), and cyanidin/peonidin-derived (D) anthocyanins on normalized total anthocyanin content. Points are sized according to normalized total anthocyanin content (square root transformed, μg/ml). Points are colored according to wavelength.

Flavonoid contents were also compared with maximal absorbance (Abs_Max_) in the ~495-525nm range, a value indicative of darkness and often used in a method for determining anthocyanin content (Supplementary Figure 1)^26^. Unsurprisingly, anthocyanin content was highly correlated with Abs_Max_ (r = 0.92, p = 2.2e-16). As with anthocyanin content, flavones were positively correlated with Abs_Max_, but the magnitude of the correlation was lower (r = 0.43, p < 2.2e-16). This may be due to the absorbance increasing effect of flavone-anthocyanin copigmentation^27^. In contrast, neither the proportion of condensed forms nor the proportion of acylated forms was significantly correlated with Abs_Max_ (p = 0.15, p = 0.07). For condensed forms this loss of a negative correlation (compared to total anthocyanin content) could be attributed to color enhancing properties of condensed forms as has been seen previously in purple corn^28^ (companion paper). The lack of a correlation between proportion of acylated anthocyanins and Abs_Max_ despite the significant relationship seen with anthocyanin content is more difficult to explain. Acylation has not been associated with a significant increase in absorbance. However, this could be an indirect effect of the correlation that exists between acylated anthocyanins and flavones in this population (Chatham and Juvik, submitted to G3). Generally, high flavone concentration was observed in lines with smaller proportions of acylated anthocyanins. These extracts could have lower acylation and lower overall anthocyanin content, but not necessarily have a correspondingly low Abs_Max_ due to the presence of flavones and the effects of copigmentation.

From a global model using all factors described above and in Supplementary Figure 1, the most parsimonious model was chosen. In this model anthocyanin content, condensed forms, flavones, and the interactions between them were all significant, but anthocyanins and condensed forms had the smallest p values and largest effect sizes by far. Variables were scaled so that effect sizes can be directly compared. The anthocyanin term had a scaled effect size of 1.5 standard deviations from the mean and a p-value too low to estimate in R. The condensed form factor had a scaled effect size of 0.5 (p = 2.7e-77) and the interaction between anthocyanin and condensed forms had an effect size of 0.6 (p = 7.3e-59). Taken together, this suggests a complex model of flux, in which each of these factors should be considered in the context of the others. Interactions between factors could directly influence Abs_Max_ as is the case for copigmentation^27^, or the interactions could be indications of the underlying biology, for example, a regulator controlling both anthocyanins and flavones.

Figure 3 shows schematics that describe two approaches to the regulation of pathway flux. With Figure 3A as a starting point illustrating how total carbon flux into the pathway is portioned between phlobaphenes, flavones, anthocyanins, and condensed forms, 3B illustrates a finite flux approach in which losses of some products are realized as gains in others (zero-sum model). Figure 3C illustrates an approach in which flux is regulated universally, where gains in one class of compounds result in gains in another (win-win scenario). The ability to breed for both scenarios simultaneously would be ideal (Figure 3D), but further manipulation of the pathway in this way will require a better understanding of genetic factors controlling flux. Flux likely has attributes of both models to varying degrees and at different places in the pathway, along with a number of other factors including environmental conditions.

**Figure 3:**
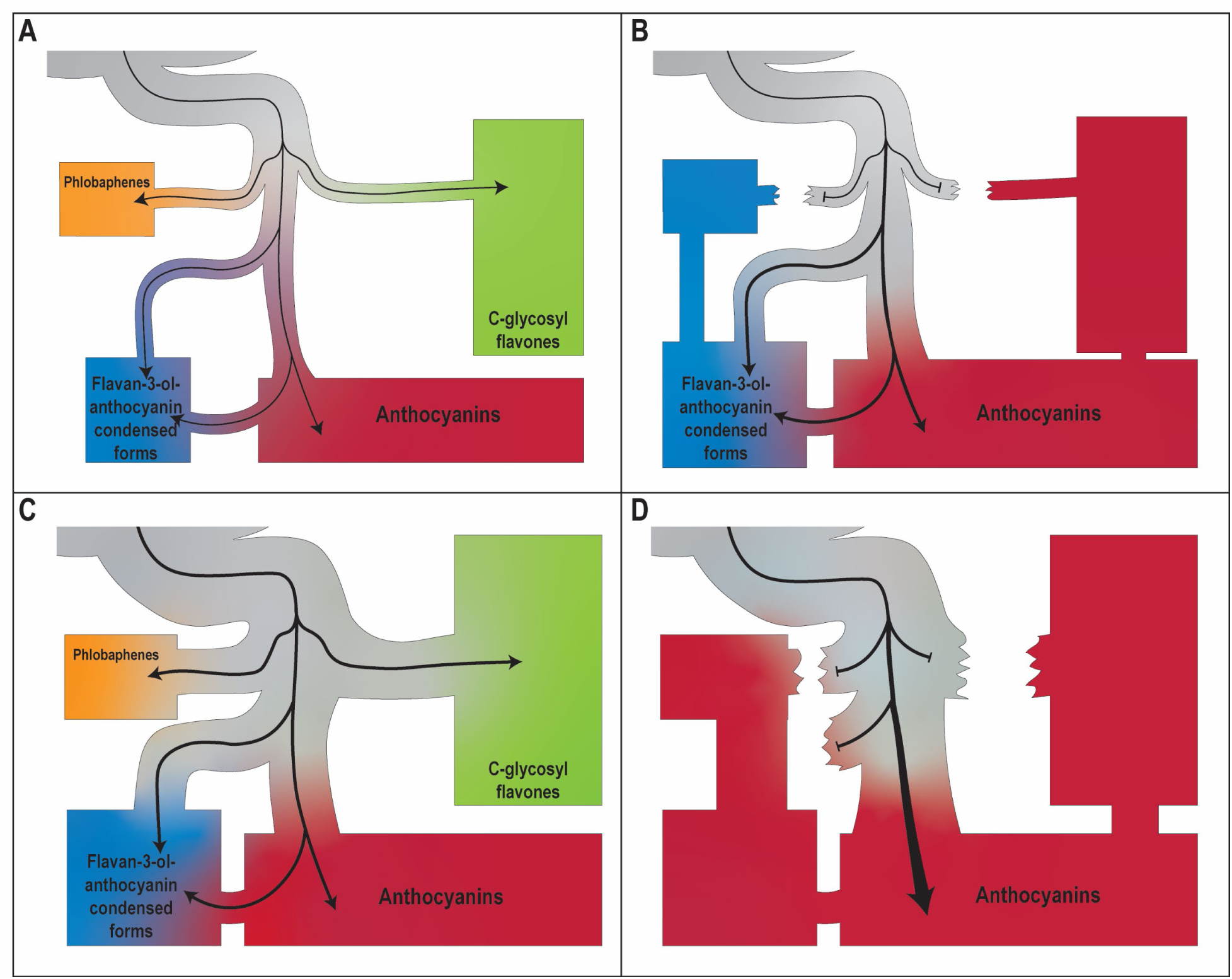
Schematic showing simplified potential strategies for increasing anthocyanin content. Diagrams are laid out to resemble the pathway presented in Figure 6.x. (A) Assumed normal flux through the pathway with all major products made. (B) Loss of structural genes or regulatory factors required for phlobaphene (orange) and flavone (green) biosynthesis (e.g. P1) shunts flux toward flavonols and anthocyanins. (C) Increased total flux through the entire pathway, and corresponding increases in all end products. (D) Combined increased flux (C) and partitioning of flux (B) toward anthocyanin biosynthesis.

### Mapping anthocyanin concentration and Abs_Max_

In mapping anthocyanin content and other flavonoid products and the subsequent selection of candidate genes, known structural and regulatory factors associated with the pathway were considered first (Table 1). A list of potential candidates based on anthocyanin and flavonoid regulatory and trafficking genes in other species was also prepared by blasting against the maize genome (Supplementary Table 1)^29^. Mapping anthocyanin content was performed using both Abs_Max_ and total anthocyanin content (Figure 4). Given the differences observed between their correlations with different flavonoid profiles, we expected slightly different mapping results. Each produced highly significant signals on Chr 1 (Chromosome 1) near *Aat1*, with the most significant SNP less than 1 Mb away. This suggests that despite being a structural gene, acylation by *Aat1* is important for achieving highly pigmented extracts. In *V. vinifera*, MATE transporters were identified that selectively transported acylated anthocyanins into the vacuole compared to their un-acylated versions^14^. While this offers a plausible explanation for a highly significant peak near *Aat1*, no anthocyanin-transporting MATEs have been identified in maize and this phenomenon has not been associated with the GST/ABC transporter system (*Bz2/Mrp3* in maize).

**Table 1:** Anthocyanin regulatory and transport / storage factors. See Chatham et al. (2019) for a complete overview of these factors.

| Locus |  | Gene product | Gene model identifier | Chromosome | Mb |
| --- | --- | --- | --- | --- | --- |
| ***Regulatory*** |  |  |  |  |  |
| *pl1* | purple plant1 | R2R3-MYB | Zm00001d037118 | Chr 6 | 113.0 |
| *c1* | colored aleurone1 |  | Zm00001d044975 | Chr 9 | 9.0 |
| *b1* | colored plant1 | bHLH (basic helix-loop-helix) | Zm00001d000236 | Chr 2 | 19.0 |
| *r1* | colored1 |  | Zm00001d026147 | Chr 10 | 139.8 |
| *hopi1* | hopi r1/b1 family member1 |  | Zm00001d026147 |  |  |
| *lc1* | red leaf color1 |  | Zm00001d026147 |  |  |
| *sn1* | scutellar node color1 |  | Zm00001d026147 |  |  |
| *pac1* | pale aleurone color1 | WDR (WD repeat protein) | Zm00001d017617 | Chr 5 | 201.8 |
| *a3* | anthocyaninless3 | unknown | no gene model |  |  |
| *p1* | pericarp color1 | MYB | Zm00001d028842 | Chr 1 | 48.4 |
| *in1* | intensifier1 | bHLH-like inhibitor | Zm00001d019170 | Chr 7 | 20.1 |
| *rif1* | r-interacting factor1 | EMSY-like / ENT domain containing protein | Zm00001d019922 | Chr 7 | 76.6 |
| ***Transport / Storage*** |  |  |  |  |  |
| *bz2* | bronze2 | GST (glutathione *S*-transferase) | Zm00001d032969 | Chr 1 | 245.0 |
| *mrp3* | multidrug resistance-associated protein3 | MRP (multidrug resistance-associated protein, ABC transporter) | Zm00001d045269 | Chr 9 | 17.5 |

**Figure 4:**
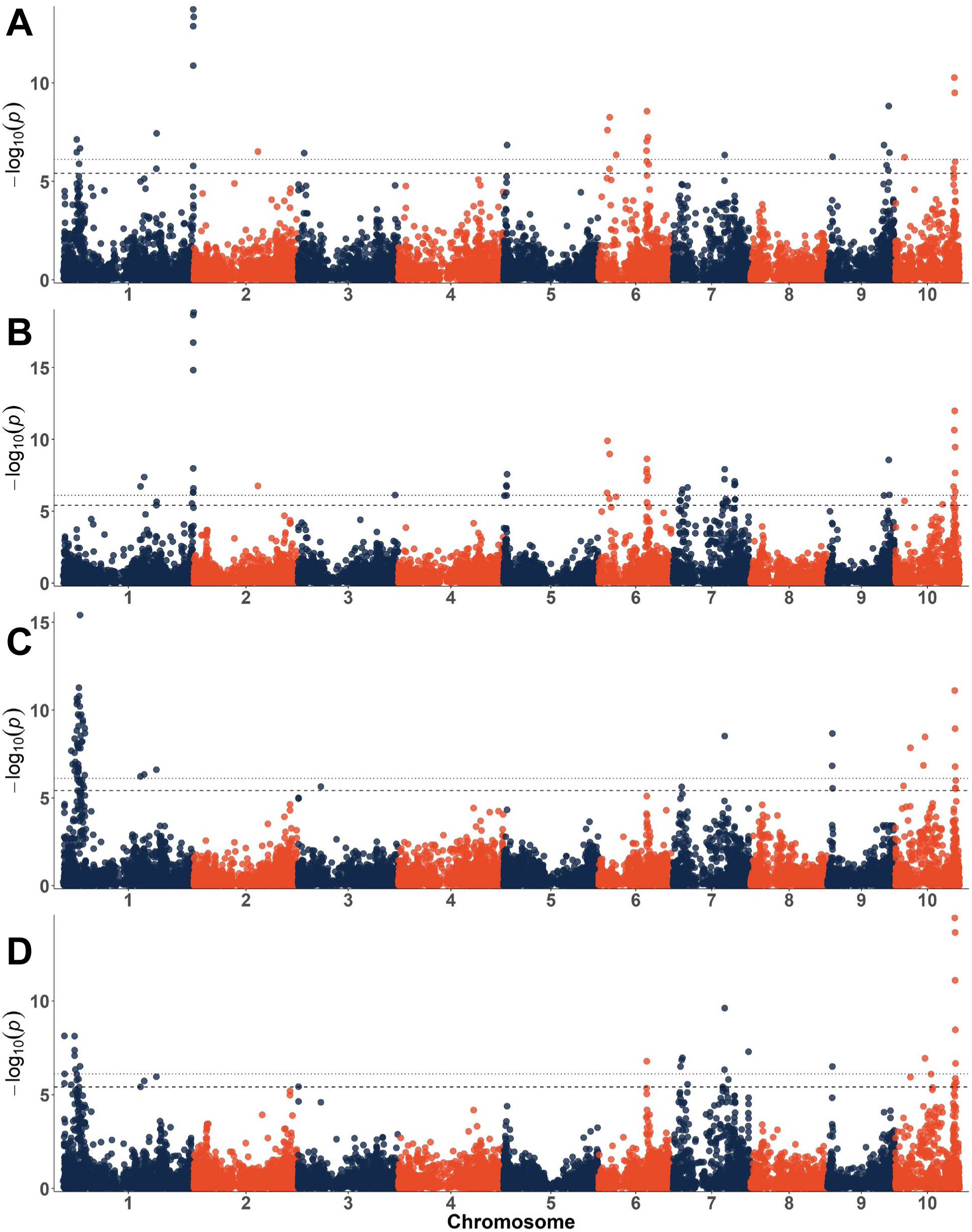
Manhattan plots for maximum absorbance (A, C) and total anthocyanin content in μg/ml (B, D). A and B contain data from the full population while C and D contain only the lines with proportions of acylated anthocyanins between 0.35 and 0.75.

Loci identified for Abs_Max_ but not for total anthocyanin content are of interest because they likely represent factors associated with increasing the darkness of extracts that are unrelated to anthocyanin content. The most plausible reason for the occurrence of these loci is involvement in flavone production and thus copigmentation with anthocyanins^27^. For Abs_Max_ a locus on Chr 1 spanning 32-40 Mb was detected and suspected to be *P1*, a major regulator of flavone biosynthesis. This signal was only observed for Abs_Max_ and not total anthocyanin content, which could be explained by the color intensifying effect of flavones in the presence of anthocyanins^27^. As a proxy for color intensity, Abs_Max_ would be expected to tag major loci associated with flavone content. Targets of *P1* were thus included as candidates in Supplementary Table 1^30^. However, the range observed is at least 8 Mb away from *P1*, located at 48 Mb. Another potential candidate considered is a gene encoding the a3 subunit of a vacuolar proton ATPase (Zm00001d028436, 34.96 Mb). If MATE transporters were found to play a role in the trafficking of anthocyanins to the vacuole, as has been observed in other anthocyanin producing species, a proton pump would likely be necessary to generate the gradient required for MATE function^2^. Individual chromosome plots showing signals and candidate gene locations can be found in Supplementary Figure 2.

Other loci observed for Abs_Max_ but not total anthocyanin content were found on Chr 3 around 14.8 Mb and Chr 9 at 11.8 Mb (Table 2). While no candidates were identified for the former, the latter is close to *Bz1* (*Bronze1*), a UDP-glucose flavonol glycosyltransferase at 11.2 Mb. Surprisingly, this signal was significant only for Abs_Max_ but not for total anthocyanin content.

**Table 2:**
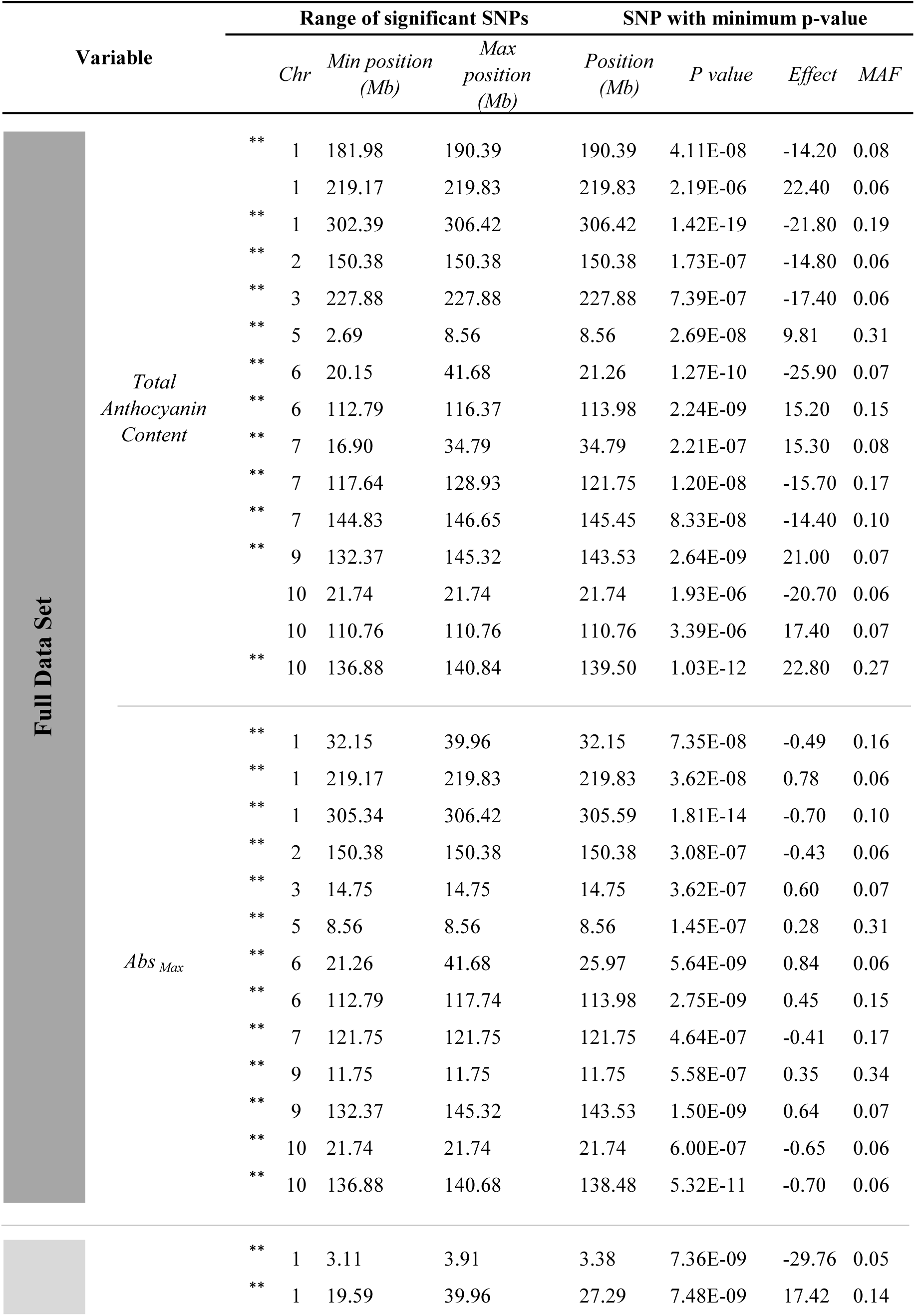
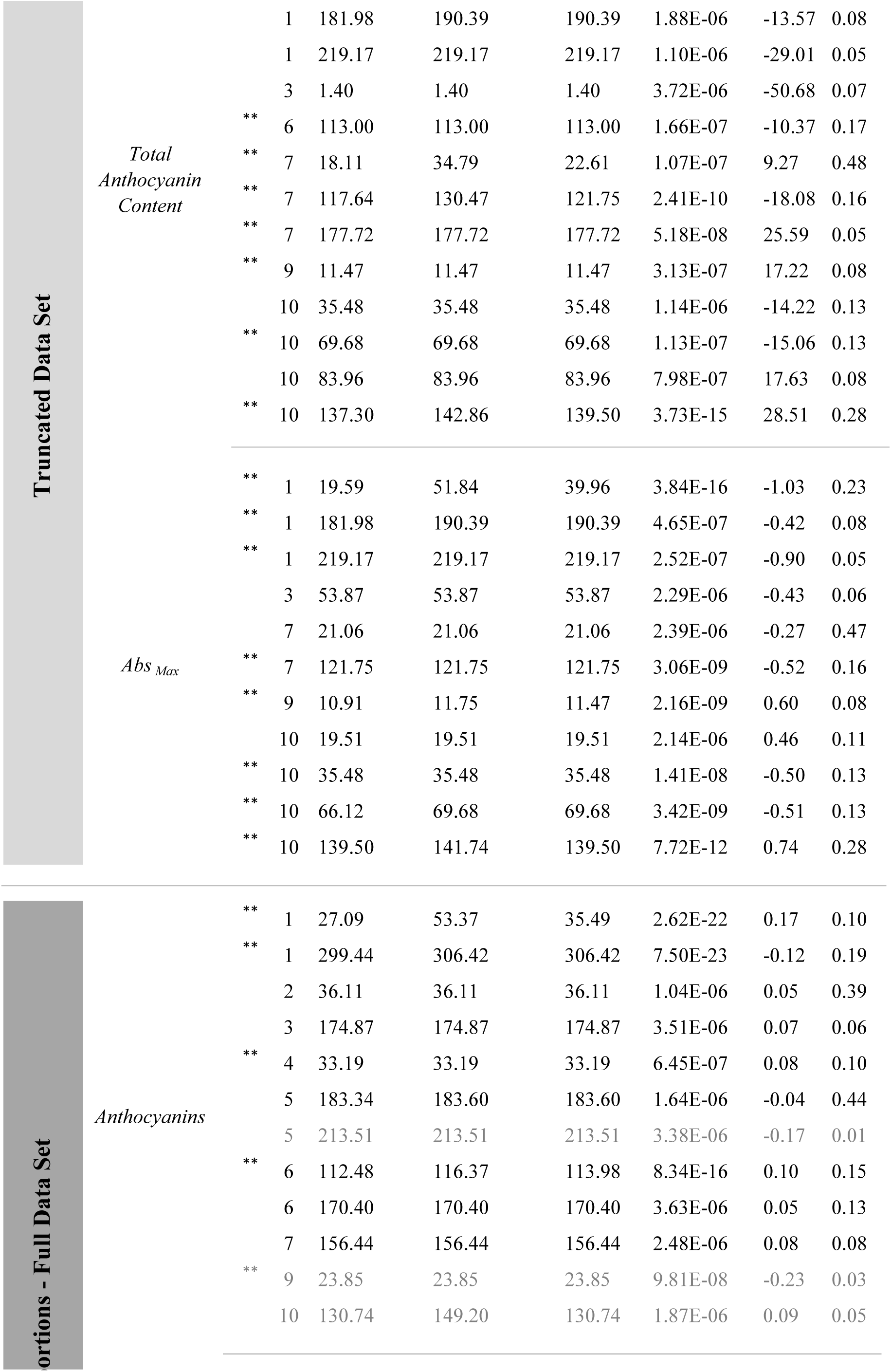
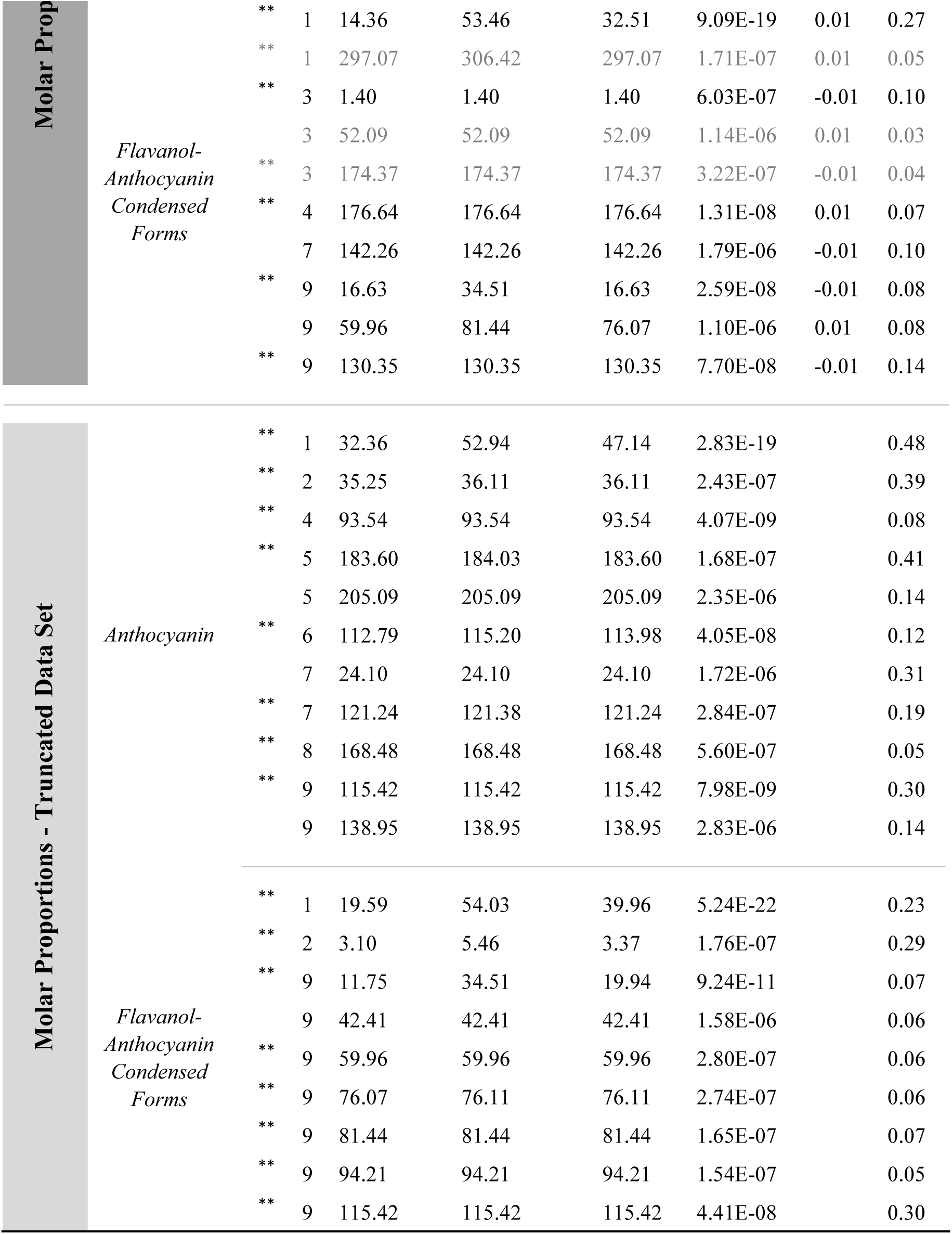
Significant SNPs^*^ for each anthocyanin variable mapped

Examining the percent of acylation in Figure 2B and Supplementary Figure 1D shows a bimodal distribution, with the lower acylation group having both fewer observations and a smaller spread of anthocyanin content and Abs_Max_. This association could be due to linkage between *Aat1* and some other factor related to anthocyanin content, but LD in the surrounding region suggests this may be unlikely. Nonetheless, to account for potential linkage, mapping was performed using acylation as a covariate, and modeling *Aat1* as a fixed factor, but the signal was still present. To overcome this, the tails of the acylation proportion distribution—less than 0.35 and greater than 0.75—were removed from the data to isolate anthocyanin content.

Using the truncated data set for mapping (acyl proportion tails removed, leaving 929 lines) eliminated the signal near *Aat1* as well as several others. Examining these eliminated loci could reveal factors associated uniquely with acylation. In addition to Chr 1 at 306.5 Mb, signals on Chr 2 at 150.4 Mb and on Chr 5 at 8.6 Mb were lost when the data was truncated, and mapping was rerun. The closest candidates identified for these two signals were a potential MATE transporter located on Chr 2 at 154.1 Mb (Zm00001d005018) and a UDP-D-glucose dehydrogenase on Chr 5 at 6.95 Mb involved in sugar metabolism and expressed in pericarps of both *P1-rr* (phlobaphene rich) and *P1-ww* (no phlobaphene) lines^30^. Another potential candidate is *Chi3* (*Chalcone isomerase3*), located at 2.6 Mb. The proximity of these signals to candidate loci can be seen in Supplementary Figure 2.

Two signals were consistently significant for all analyses. The first, on Chr 1 at 219.8 Mb, did not have any immediately obvious candidates. A signal in this area was also identified previously for several traits in this article’s companion (Chatham and Juvik, submitted to G3). The significant SNP for this signal is about 4 Mb away from *myb83*, discussed previously, and is very close to a WRKY transcription factor located at 219.8 Mb (wrky25; Zm00001d032265). WRKY transcription factors are a more recent addition to the anthocyanin regulatory complex identified in both *Arabidopsis* (*TTG2*) and Petunia (*PH3*), and these likely interact with the WD repeat protein component of the MBW regulatory complex^3^. The candidate WRKY, wrky25 is also highly expressed in pericarp tissue^31^; however, when *TTG2* and *PH3* were blasted against the maize genome, wrky25 was a weak match. We also identified a wrky candidate (wrky62, Zm00001d035323) near a QTL on Chr 6 for anthocyanin content, but it too was a weak match for both TTG2 and PH3.

With what is known regarding the regulation of anthocyanin content in pericarp, and previous reports breeding for purple corn^32,33^, we expected to find *B1* and *Pl1*. Instead, SNPs proximal to *R1* (less than 1 Mb away) were found in all four analyses. The *R1* locus is diverse, and while usually associated with aleurone color, some alleles of *R1* (e.g. *R1-ch*), and its neighboring (or allelic) loci *Sn1* (*Scutellar node1*)^34^, *Lc1* (*Leaf color1)*^35^, and *Hopi*^36^ can produce anthocyanins in pericarp^6^. Additional research will be required to identify the specific bHLH allele in AR and whether it interacts with other proteins in the manner described previously for aleurone-pigmented lines.

*R1* is a bHLH transcription factor and part of the MBW ternary regulatory complex associated with anthocyanin biosynthesis across species^3^. Therefore, we expected to also find a MYB and a WD40 as well. While not as obvious as for *R1*, a peak can be seen near the plant color associated MYB, *Pl1* (Chr 6 at 113.0 Mb), in Figure 4, but no signal was found near *Pac1* (Chr 5 at 201.8 Mb), the WD repeat protein component of the MBW regulatory complex. *R1* has also been shown to interact with *Rif1* (*R-interacting factor1*), potentially to regulate control of early versus late biosynthetic steps in the anthocyanin pathway^5^. No signal was observed close to *Rif1* (Chr 7 at 76.6 Mb) either, though it is possible that some key components of the regulatory complex (e.g. *Rif1* and *Pac1*) are fixed in the Apache Red population, offering an explanation for these missing signals. Alternatively, pericarp-specific alleles of these could exist that have not yet been identified.

In addition to those described above, several signals were identified that were present in some but not all the described mapping approaches (Figure 4). On Chr 1, two significant SNPs were found at 182 and 190 Mb, for which the closest candidate found was a set of potential ANR/LAR like proteins (Zm00001d031488, Zm00001d031489). On Chr 7 we identified a peak near *In1*, a recessive intensifier of anthocyanin biosynthesis in aleurone-pigmented lines (Supplementary Figure 2). To our knowledge, there have been no reports of *In1* functioning in pericarp. However given the capacity of *in1* to significantly increase anthocyanin content in aleurone^37^, this should be explored further as a potential strategy to maximize anthocyanin content in pericarp. Another peak on Chr 7 was found close to myb162 (Supplementary Figure 2).

### Mapping molar flux

While the mapping variables used up to this point primarily correspond to the model of increased overall flux from Figure 3C, we also wanted to look for regulators that may fit the competitive flux model seen in Figure 3B. To do this, concentrations of individual anthocyanin and flavone compounds were converted from μg/ml to mols/ml, and the proportion of the total molar flux into the pathway was calculated and mapped for anthocyanins, flavones, and condensed forms (Figure 5). Given the relatively small proportions of condensed forms, anthocyanin and flavone proportion made up the majority of flux through the pathway for these calculations and thus mapping both anthocyanin and flavone proportions was redundant (e.g. flavone and anthocyanin proportions are tightly inversely correlated). Manhattan plots from flavone flux and anthocyanin flux were nearly identical and the same signals were observed for both. Nonetheless, these variables still represent partitioning between branches of the pathway, and candidates identified may play key roles in regulating this.

**Figure 5:**
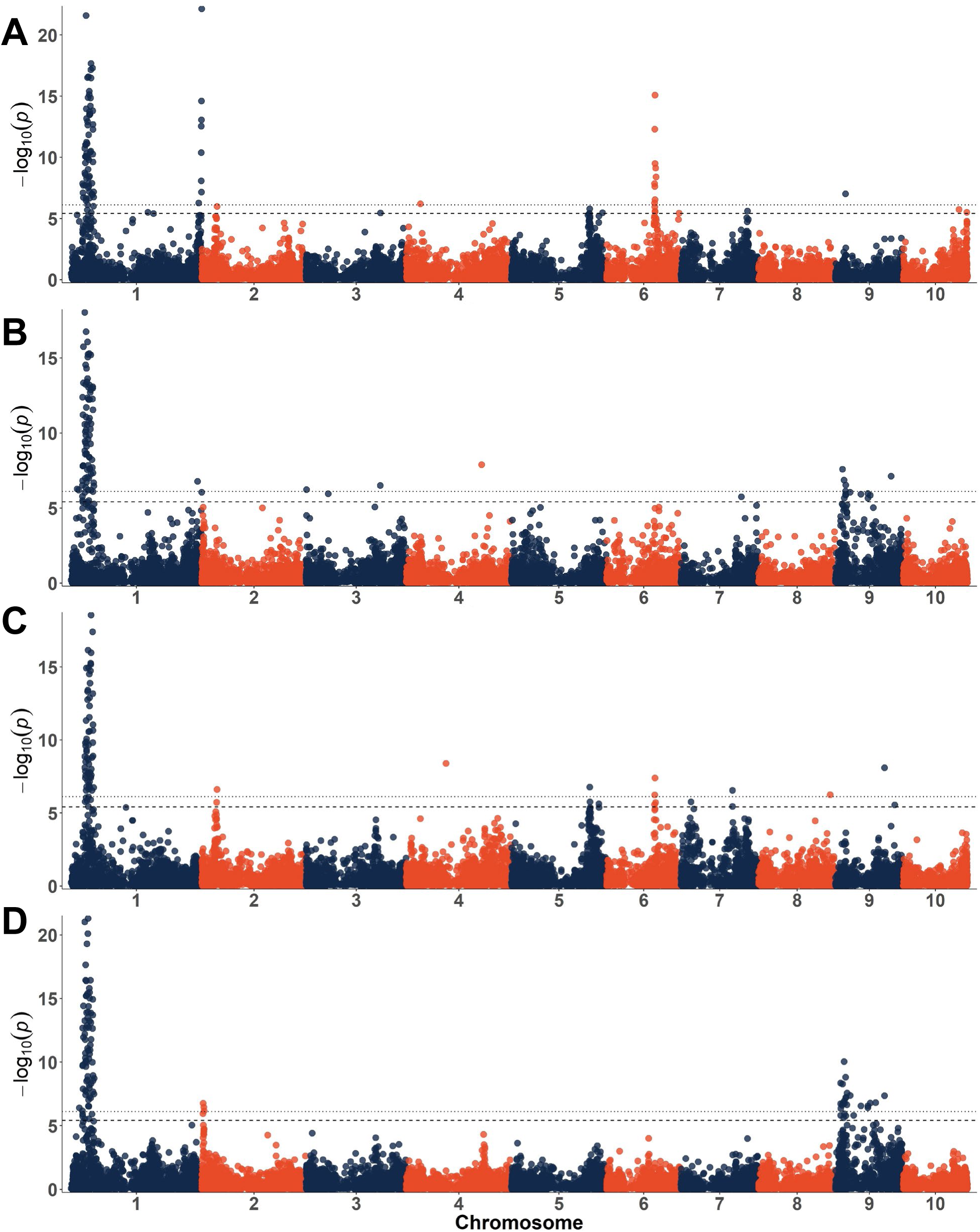
Manhattan plots for the partitioning of flux through the flavonoid pathway, showing proportion anthocyanins and the (B) proportion of condensed forms

For all proportions mapped, signals close to *P1* and *Aat1* were identified. *Pl1*, the bHLH regulatory factor identified previously, was conspicuous both for proportions of flavone and anthocyanin flux. While *R1* showed up consistently when mapping anthocyanin content overall, it was barely detectable when looking at proportions of flux through the pathway. This suggests *Pl1* may play a more significant role in the competition for flux through the pathway, influencing the ratio of flavones and anthocyanins, while *R1* may contribute more to overall flux.

Similar to results of mapping absorbance, the peak at the beginning of Chr 1 was close to *P1* but not nearly as close as other candidates have been to their assumed loci (e.g. *Pr1, R1*). The most significant SNP was more than 10 Mb away from *P1*. This could be explained if large linkage blocks in this area were detected. Pairwise recombination frequency was calculated for the region on Chr 1 between 33 and 50 Mb (Supplementary Figure 3). *P1* is located at 48 Mb, but the most significant SNP for anthocyanin/flavone flux was found at 35 Mb. Higher linkage in these areas could contribute to the gap between the most significant SNPs and *P1* (Supplementary Figure 4). The frequency with which we found *P1* for various traits, both here and in this article’s companion, is somewhat puzzling given the degree of selective pressure for anthocyanin content during the development of the AR population (Figure 1). One explanation may be the presence of multiple loci located in a relatively small area, making separation difficult. When this region was searched, a candidate [Zm00001d028436 (vacuolar proton ATPase a3)], at 35 Mb was identified. This or other additional candidates could explain the frequent detection of the peak on Chr 1 in phenotypes not expected to be influenced by *P1*. However, it cannot be determined without further research whether the positive correlation between flavones and anthocyanins is a cause (leading to detection of *P1* in anthocyanin mapping) or an effect (correlation resulting from linked anthocyanin and flavone loci) of the significant peak observed on Chr 1.

We also identified a signal near *Pr1* when mapping the proportion of anthocyanin and flavone content (Figure 5). In general, we observed that *pr1/pr1* lines typically had higher flavone content compared to *Pr1/-*lines. Moreover, the only flavone compounds detected here in abundance were apigenin derived (monohydroxylated B-ring), consistent with their presence in *pr1/pr1* lines containing primarily pelargonidin (also monohydroxylated B-ring). Therefore, we suspect the occurrence of the signal on Chr 5 is an artifact of population structure. Yet interestingly, studying *pr1* in the context of C-glycosyl flavone production showed that recessive *pr1* increased apimaysin content (monohydroxylated) but *Pr1* had little to no effect on maysin content (dehydroxylated B-ring) suggesting the presence of divergent pathways and the expression of an additional F3’H^38^. This might explain the lack of luteolin-derived flavones (dehydroxylated B-ring), even in the presence of *Pr1* (cyanidin-dominant lines). Regarding anthocyanin content, a structural gene like *Pr1* seems unlikely to influence total anthocyanin content yet we have noticed that pelargonidin lines generally have lower ceilings of anthocyanin production in comparison with cyanidin-dominant lines in the purple corn germplasm^39^. Additional research would be required to determine whether there is a true biological explanation, such as substrate specificity, or if this is an artifact of population structure.

Several other significant SNPs or SNP ranges were observed (Table 2). On Chr 7, a peak was found at 156.4 Mb, which aligns closely with a potential ortholog of the *Arabidopsis TT12 (Transparent Testa12*) MATE transporter of flavonoids (Zm00001d021629, 158.6 Mb)^12^ and other flavonoid transporting MATEs in other species (Supplementary Figure 3)^14,15,17^. This is particularly interesting given the highly significant effect of SNPs near *Aat1* for both competitive flux and overall flux (Figures 4 and 5)*. Vv*AM1, *Vv*AM3, and *Mt*MATE2 all preferentially transport acylated anthocyanins ^14,15^, while MtMATE1 is specific to epicatechin^15,40^. A candidate MATE with specificity for acylated anthocyanins could explain how *Aat1* appears to contribute so significantly to anthocyanin content. A phylogeny of candidate MATEs and MATEs associated with flavonoid transport in other species can be found in Supplementary Figure 5. Differences in specificity could also play a role in the relative amounts of condensed forms versus anthocyanins that are stored in the vacuole. We also found a single significant SNP on Chr 9 (23.9 Mb), for which the closest candidate we found is another MYB transcription factor (myb148, 26.7 Mb; Supplementary Figure 2). While this QTN was highly significant, minor allele frequency was low (0.03) raising concern that this could be a false positive result.

While in this article’s companion, total condensed forms and the proportion of condensed anthocyanins were mapped, here we mapped the molar proportion of condensed forms synthesized compared to total pathway flux. The signal on Chr 9 (130.67 Mb) is suspected to correspond with *Fns1* (Chr 9 at 30.7 Mb). While *Fns1* is responsible for the formation of flavones, it could participate in the competition of flux toward flavones versus anthocyanins and flavan-3-ols, thereby influencing condensed forms indirectly. *Fns1* is regulated by both *P1* and the MYB (*Pl1/C1)* and bHLH (*B1/R1*) system, which could contribute to the frequent co-occurrence of anthocyanins, flavones, and condensed forms in Apache Red extracts^30,41^.

A signal on Chr 3 at 174.4 Mb was also identified 1.3 Mb from a candidate ANR (anthocyanidin reductase) (Zm00001d042594). In this articles companion paper, several potential ANR/ LAR candidates were proposed, but no signals were produced close to this particular candidate. Candidates identified previously clustered with LARs while this candidate groups with ANRs. While LARs have only been shown to produce 2,3-*trans* flavan-3-ols (e.g. catechin) and ANR shown primarily to produce 2,3-*cis* flavan-3-ols (e.g. epicatechin), epimerase activity converting 2,3-*cis* flavan-3-ols to 2,3-*trans* flavan-3-ols has been found in several species, suggesting an ANR may be sufficient for the production of both flavan-3-ol types^42–44^. However, LARs have also been implicated in proanthocyanidin biosynthesis by regulating the number of epicatechin starter and extension units used to create polymers^45^. This has not been established for the formation of condensed forms, but a similarly functioning LAR could play a role in the dimerization of flavan-3-ols and anthocyanins to make condensed forms^2^.

Mapping flux toward condensed forms produced several signals that had not been identified in previous analyses, including a hit on Chr 4 at 176.6 Mb. A UDP-D-glucuronic acid 4-epimerase (Zm00001d052053) regulated by *P1*^30^ was found 1.2 Mb away, and putative MATE transporter (Zm00001d051886) with similarity to *Vv*AMT anthocyanin-transporting MATEs was found 3 Mb away from the peak on Chr 4. However the closest candidate (<0.2 Mb) is an ATPase (Zm00001d052022) similar to TT13 from *Arabidopsis*, a P-type H^^+^^ ATPase required for epicatechin transport into the vacuole and proanthocyanin accumulation in seeds. TT13 provides the H^+^ gradient required to fuel TT12, a MATE transporter of epicatechin in *Arabidopsis*^46^. This candidate also has similarity to *PH5* from petunia, an H^+^ P-ATPase that controls vacuolar pH and flower color^47^.

We also observed several signals that had been previously detected when mapping other traits. This includes a peak at the beginning of Chr 3 near *tcptf33*, a blast hit for *TCP3* from *Arabidopsis* which interacts with the MBW regulatory complex in the control of flavonoid biosynthesis^48^. This same signal was observed when mapping total condensed forms in this articles companion paper (Chatham and Juvik, submitted to G3). Similarly, signals on Chr 3 around 52 Mb and on Chr 7 at 142 Mb were again detected. Potential candidates for these include myb40 (Chr 3, 54.36 Mb) and a potential LAR/ANR ortholog (Zm00001d020970, Chr 7, 138.21 Mb) or glucosyltransferases (Zm00001d021167, Zm00001d021168, Chr 7, 144.7 Mb). Several potential MYBs have been identified here and in this article’s companion. A phylogeny of known anthocyanin activators and repressors from other species and maize candidates can be found in Supplementary Figure 6.

**Figure 6:**
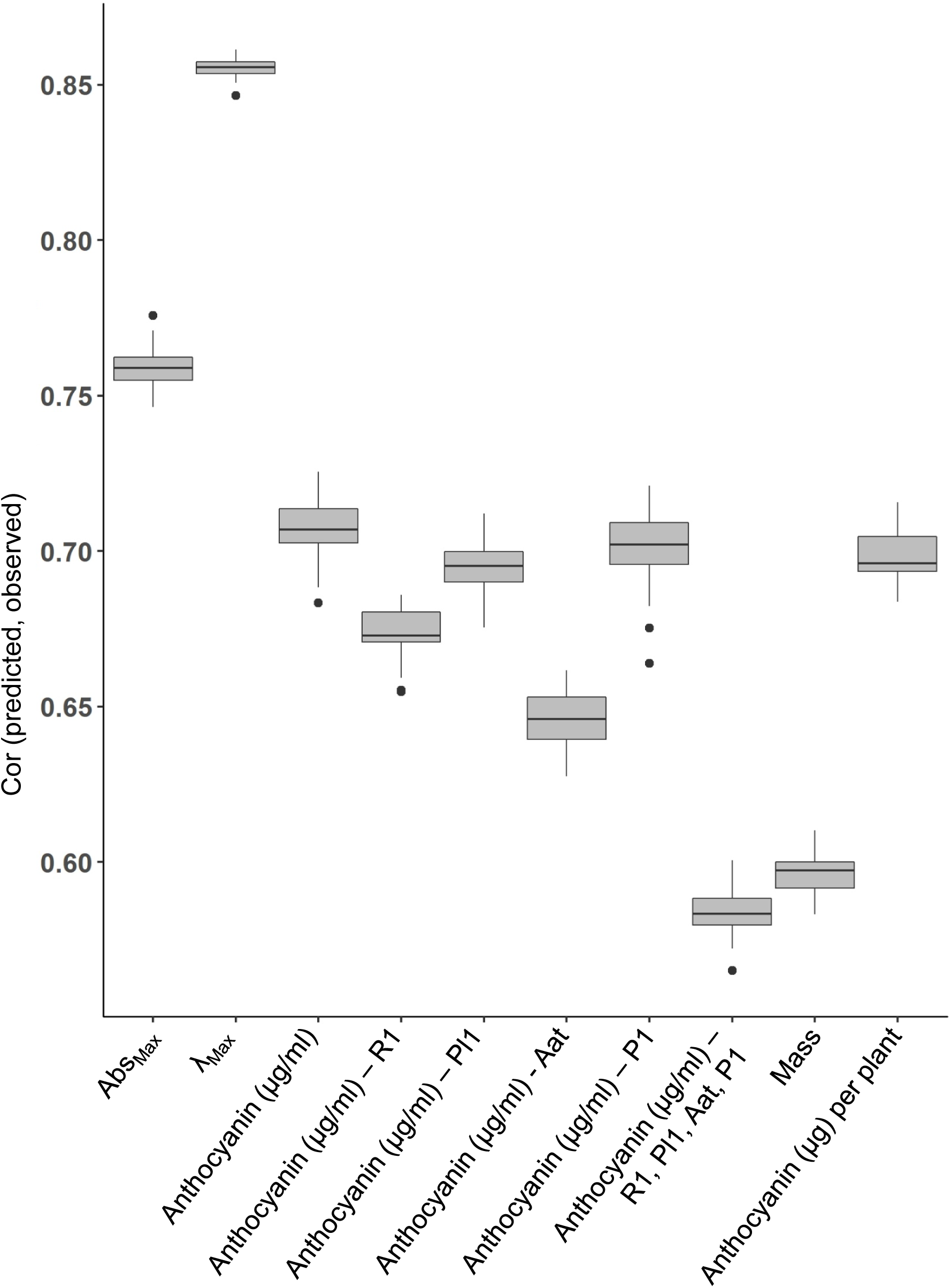
Prediction accuracy of genomic selection.

While a single significant SNP at the beginning of Chr 9 was found for anthocyanin proportions, a large peak with a string of significant SNPs was found here for proportions of condensed forms, with several trailing significant SNPs extending toward the centromeric region (Supplementary Figure 3). While the most significant SNP for this peak was located at 16.6 Mb, the width of the region makes identifying candidates difficult. Previously, this same region at the beginning of Chr 9 has been identified in association with flavonoid content. *Bp1* (*Brown pericarp1*) is located in bin 9.02 (11.8 – 23.3 Mb in B73 RefGen_v3)) and is a locus associated with modifying red pericarp to brown when recessive and in the presence of *P1*^49^. It was therefore suggested to play a role in the formation of phlobaphenes^50^. This same area was later associated with maysin biosynthesis and corn earworm antibiosis (*recessive enhancer of maysin1 (rem1*)) and was mapped between *Bz1* (11.2 Mb) and *Wx1* (23.1 Mb)^38^. In addition to phoblaphenes and C-glycosyl flavones, we have now found this region to be important for the proportion of flavonol-anthocyanin condensed forms produced. Interestingly, this QTL region was not nearly as evident for the proportions of anthocyanin / flavone content, which based on previous reports would have been expected. Nor did we identify this same region for total C-glycosyl flavone content in this article’s companion. Several scenarios could account for the significance of this region in multiple traits. Proximally located genes with different functions and their resulting linkage could help explain the correlation between flavones and condensed forms. Alternatively, the *rem1 / bp1* region might contain a single gene whose product acts at a point in the phenylpropanoid pathway prior to differentiation between phlobaphene, flavone, and flavanol branches. Similarly a single gene in this region could act as a regulator at multiple points in the pathway.

Given this region’s involvement in phlobaphene, flavone, and condensed form content, several potential candidates were considered. A plasma membrane ATPase (Zm00001d045122, 13.45 Mb), a glucosyltransferase labeled as an anthocyanidin 5,3-O glucosyltransferase (ufgt3, Zm00001d045254, 17.2 Mb) and *ZmMrp3* (17.51 Mb) the ABC transporter involved in anthocyanin transport^11^.Locations of these are shown in Supplementary Figure 3. Condensed forms and C-glycosyl flavones are sequestered in the vacuole, while phlobaphene polymers are destined for the cell wall. If a MATE transporter were required to transport flavones and flavan-3-ols into the vacuole (as in *Arabidopsis*), and to move phlobaphene precursors across the cell membrane, a generalized ATPase providing the necessary proton gradient could play a role in the transport of all three classes of compounds. Failure to transport the necessary precursors could result in compound degradation leading to brown kernels lacking phlobaphenes and a decrease in maysin content. Similarly, *Mrp3*, while associated with anthocyanin transport into the vacuole, could function similarly for other compounds. In this case, *mrp3* could result in degraded phlobaphene precursors, flavones, and flavan-3-ols.

The glucosyltransferase candidate could also be versatile enough to play a role in the accumulation of flavones, flavan-3-ols, and flavan-4-ols. Anthocyanidins must be glycosylated for stabilization and transfer to the vacuole, and thus failure of the anthocyanin glycosyltransferase *Bz1* in aleurone results in the degradation of anthocyanidins and bronze kernels instead of blue, purple, or pink kernels (depending on the anthocyanin makeup)^1^. While little is known about the transport and storage of C-glycosyl flavones or the transport, polymerization and storage of phlobaphenes, an analogous scenario is plausible in which flavan-4-ols (phlobaphene precursors), flavan-3-ols, and flavones are degraded before reaching their storage destination. More evidence for this hypothesis comes from the phenotype described for the *Bp1* locus (red kernels become brown in the presence of *P1* and *bp1/bp1)*, which is akin to the *bz1* mutant phenotype (blue/pink/purple kernels to bronze in the presence of *bz1/bz1*). However, flavones are C-glycosylated, flavan-3-ol components of condensed forms are not known to be glycosylated at all before condensation with an anthocyanin, and the pathway for phlobaphenes is unknown after the formation of flavan-4-ols. Furthermore, a bifunctional C-/O-glycosyltranferase associated with C-glycosyl flavone biosynthesis and utilizing both apigenin and luteolin as substrates has already been identified on Chr 6^51^. However, this candidate is supported by our observations of condensed form species. Condensed forms from AR lines consist of an (epi)catechin or (epi)afzelechin linked to an anthocyanidin 3,5-diglucoside. Interestingly we have not found free (non-condensed, no flavonol unit) anthocyanidin 3,5-diglucosides nor have we found condensed forms with anthocyanins containing a single glucose moiety^52^. This supports the ufgt3 candidate, especially given the rise in significance seen when mapping the proportion of condensed forms^53^.

Given the strong signal observed near *Aat1* for both of the above mapped traits, lines with proportions of acylated anthocyanins in the tails of the distribution were removed from the data, as described above. This effectively eliminated the signal near *Aat1.* Signals near *P1* on Chr 1 were still observed, and for anthocyanin proportions the most significant SNP was much closer to *P1* than observed previously (47.14 Mb). Several new signals also emerged for both analyses.

For anthocyanins (and flavones), the most obvious differences between the full and truncated data set were new signals that emerged on Chrs 7 and 9, however several other new signals were also present (Table 2). For Chr 7, signals at 24.1 Mb and 121.2 Mb were found. For the former, the closest candidates were *In1* (20.1 Mb) and *UFGT2* (UDP-flavonol-glycosyltransferase2), a glycosyltransferase (Zm00001d019256, 24.2 Mb), while for the latter the closest candidates were 4-5 Mb away (predicted flavonoid 3’-monooxygenase (Zm00001d020628) at 125.5 Mb and myb162 at 115.7 Mb. No candidates were found for the new signals on Chr 9. The peak on Chr 5 near *pr1* was still evident, but several significant SNPs were found around 205 Mb. The nearest candidates are a LAR/ANR candidate (Zm00001d017771) at 206.3 Mb and *Pac1* at 201.8 Mb.

When mapping condensed forms with the truncated data set a peak at the beginning of Chr 2 was found less than 1 Mb from *Fht1* (3.56 Mb), the flavanone 3-hydroxylase associated with the conversion of flavanones to dihydroflavanols. This step is required for the formation of both flavan-3-ols and anthocyanins; but surprisingly, this peak was only observed when mapping condensed forms. Signals on Chr 9 were also observed across a wide range as discussed previously for the un-truncated data set.

### Genomic selection

While understanding flux through the anthocyanin pathway may be beneficial for identifying large effect loci to be used in a marker-assisted selection (MAS) type approach, we also wanted to test the efficacy of a genomic selection approach. While MAS relies on a small number of large effect loci, the type easily detected using association mapping studies, genomic selection uses many small effect loci to predict a genomic estimated breeding value (GEBV). Therefore, complex traits with many small effect loci may be better suited to a genomic selection approach. The results for mapping proportions of flux through the pathway were fairly straightforward, with only a handful of significant loci detected. However, mapping the magnitude of flux was complex and many significant loci were identified. Genomic selection may be a better strategy for breeding this type of complex quantitative trait^54^.

Using five-fold cross validation we tested the effectiveness of genomic selection for several different traits (Figure 6). Focusing first on visual qualities, absorbance (Abs_Max_) and wavelength (λ_Max_), genomic selection was highly effective, with prediction accuracy averages of 0.76 (Abs_Max_) and 0.86 (λ_Max_). Accuracy was the lowest for mass (g/ear; 0.60) unsurprisingly. However, given that the ears were self-pollinations made by hand, there is likely variability in mass due to differences in pollination efficiency in addition to biological variation. Mass ranged from 4-136 g per ear. Nonetheless this prediction accuracy is relatively high for a trait as complex as grain yield. This is likely attributable to the relatedness between samples, all having been generated from a single landrace^55^. The prediction accuracy average for anthocyanin content was 0.71, but we also calculated anthocyanin content on a per-plant basis, using anthocyanin concentration in μg/g of whole corn kernels and multiplying by grain yield. Ultimately, the feasibility of using corn for a natural colorant depends on the economical production of anthocyanins. This is influenced by both grain yield and grain anthocyanin content. When five-fold cross validation was performed for this trait, we again saw high prediction accuracies, with a mean of 0.70. This suggests a genomic selection approach may be highly effective in breeding for anthocyanin yield in the AR population.

Considering the models illustrated in Figure 3, we hypothesize that a dual approach selecting for both increased overall flux and increased partitioning toward anthocyanin biosynthesis could have a dramatic effect on anthocyanin content. Given the complexity observed with overall flux and the relatively few number of loci associated with partitioning of flux, we attempted to predict breeding value by adding known factors associated with flux to the genomic selection model as fixed effects. This approach has been suggested to improve prediction accuracy^56^. We added *R1, Pl1, P1*, and *Aat1* each individually to the GS model, but prediction accuracy did not improve from the 0.71 value observed for the GS model alone. We also added all of these to the model simultaneously which significantly reduced accuracy. These findings are similar to those recently published in a study simulating the addition of fixed effects in this manner^57^.

Together this data suggests that if inputs (cost, time, etc.) are equal, the GS approach is likely to outperform a MAS approach in breeding for anthocyanin content within the AR landrace. However, this accuracy is likely to decrease when parent offspring validations are calculated outside of the training population and if GS is used for making selections once new germplasm is introduced. In this scenario, a training set would need to be defined that more accurately represents the remainder of the population to be tested. While GS may be a faster way to breed for maximal anthocyanin content, it can underestimate the effect sizes of large effect loci. Furthermore, understanding the players involved in regulation and transport of anthocyanins may provide insight into the general biological processes surrounding metabolite trafficking. Identifying and understanding loci with large effects on the phenotype may also be useful if gene editing were to be used for increasing anthocyanin content.

### Conclusion

Highly concentrated purple pericarp is only found in a select portion of maize germplasm, most of which are not adapted to a Midwest environment and do not have yields comparable to today’s hybrids. Apache Red, a pericarp-pigmented purple corn variety, offers high anthocyanin concentration and variability in extract colors^52^. However, due to linkage drag from donor purple corn parents, many generations of backcrossing to elite, adapted inbreds (as the recurrent parent) may be required to recover desired agronomic traits while maintaining anthocyanin concentrations. Furthermore, as different desirable traits are identified from the donor, (i.e. different anthocyanin types / decorations), additional introgression pipelines may be needed.

Pre-breeding of landraces or other un-adapted germplasm however, could make desirable traits from these sources more accessible by helping to eliminate deleterious alleles that may later be “dragged” via linkage during backcrossing^58^. Figure 7 illustrates the benefits of a landrace pre-breeding scheme in which the number of required backcross generations is reduced compared to an approach using a direct landrace by elite parent cross (Figure 7A). The high prediction accuracies observed here for genomic selection show that significant improvement could be made within landrace populations. Moreover, knowledge of the key factors associated with maximizing anthocyanin content could help ensure their fixation in the development of superior donor parents. This could improve the efficiency of introgression, maximizing the transfer of desirable traits while reducing movement of those that are undesirable. Lastly, superior donor parents could streamline the introgression process as breeding goals shift or change, saving the breeder from having to return to un-adapted sources.

**Figure 7:**
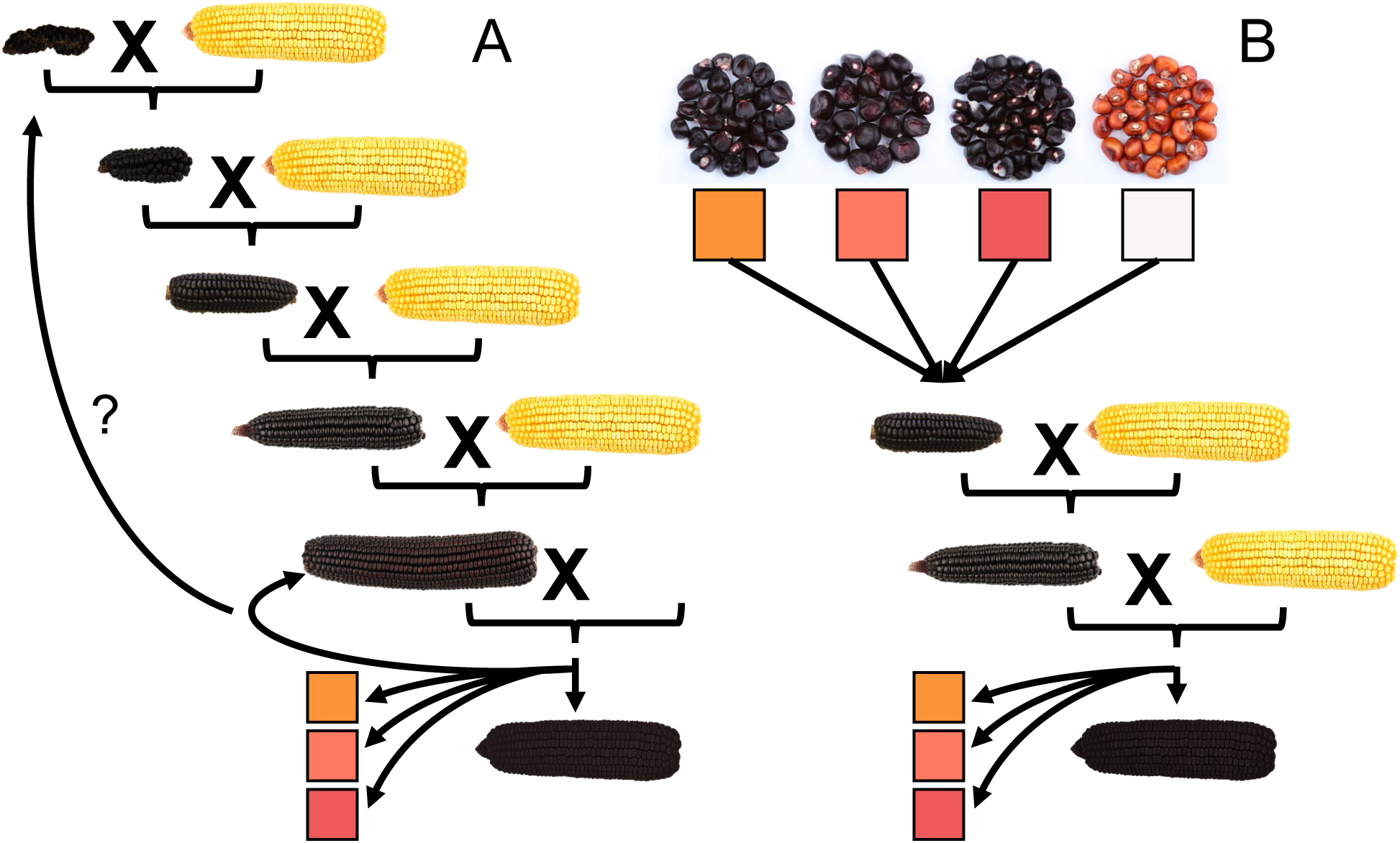
Traditional introgression (A) compared to landrace improvement (B). When multiple traits are required from a landrace, improvement of landrace materials prior to introgression may reduce linkage drag and the number of backcrosses required to regain necessary agronomic traits while maintaining the landrace sourced trait.

## List of Abbreviations Used

ABC: ATP-Binding Cassette
AR: Apache Red
bHLH: basic Helix-Loop-Helix
GBS: Genotyping-By-Sequencing
HPLC: High-Performance Liquid Chromatography
MAF: Minor Allele Frequency
MATE: Multidrug and toxic compound extrusion
MYB: derived from “Myeloblastosis protein”
PCA: Principal Component Analysis
SNP: Single Nucleotide Polymorphism
QTL: Quantitative Trait Loci

## Declarations

### Ethics Approval

Not applicable

### Consent for Publication

Not applicable

### Availability of Data and Material

All genotype and phenotype datasets will be made available at the time of publication via Figshare under project name “Linking anthocyanin diversity, hue, and genetics in purple corn”. (Private links for review: https://figshare.com/s/e116278ff976d2e19c6e and https://figshare.com/s/18d7f81d7d2e9b1c47b1)

### Author’s Contributions

LAC developed, genotyped, and phenotyped the Apache Red population, analyzed the data, and prepared the manuscript. JAJ assisted in conceiving the study and revising the manuscript. All authors read and approved the manuscript.

## Acknowledgements

We would like to thank Dr. Pat Brown for assisting with GBS library construction.

---

**Supplementary Table 1.**
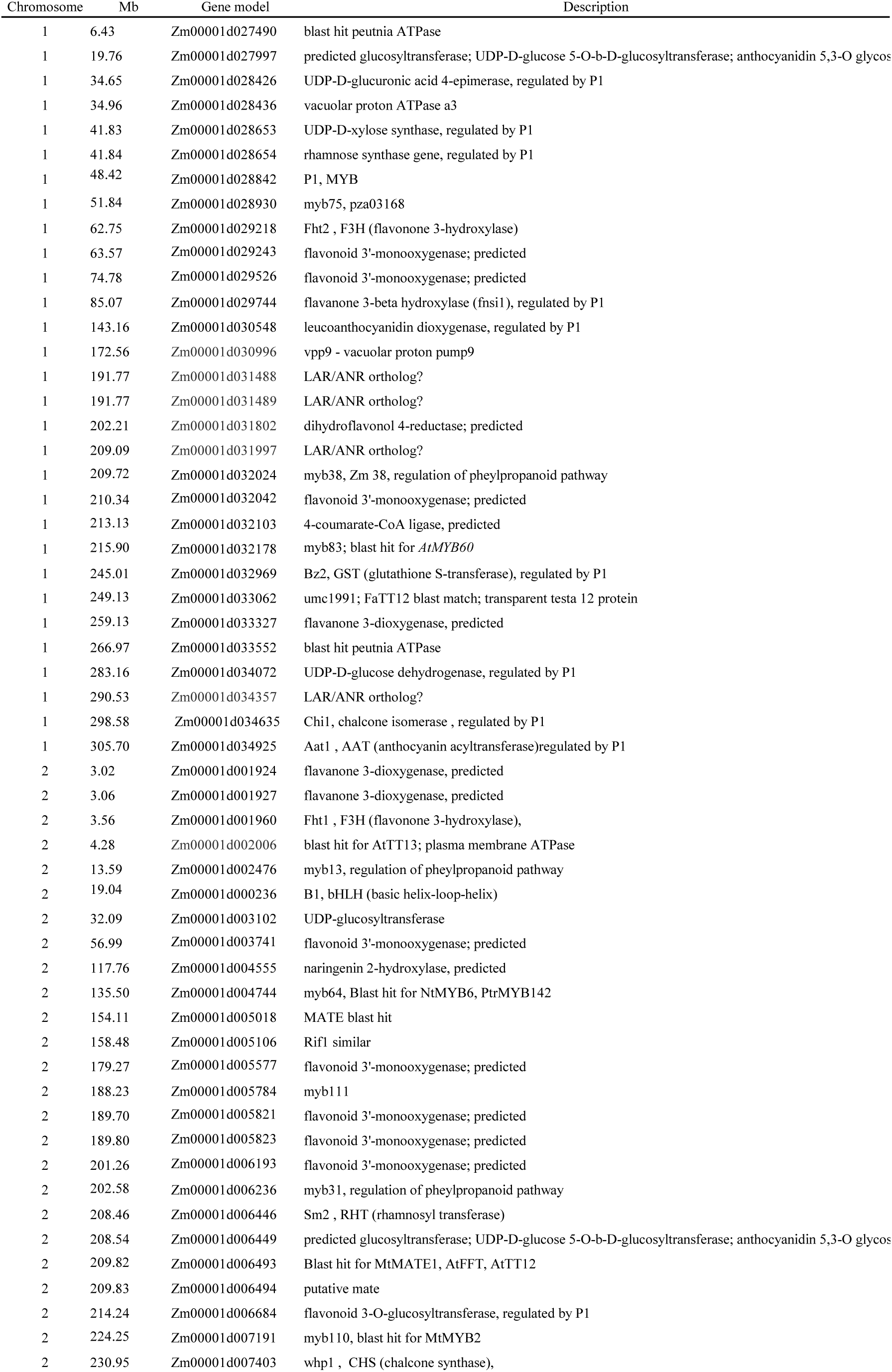
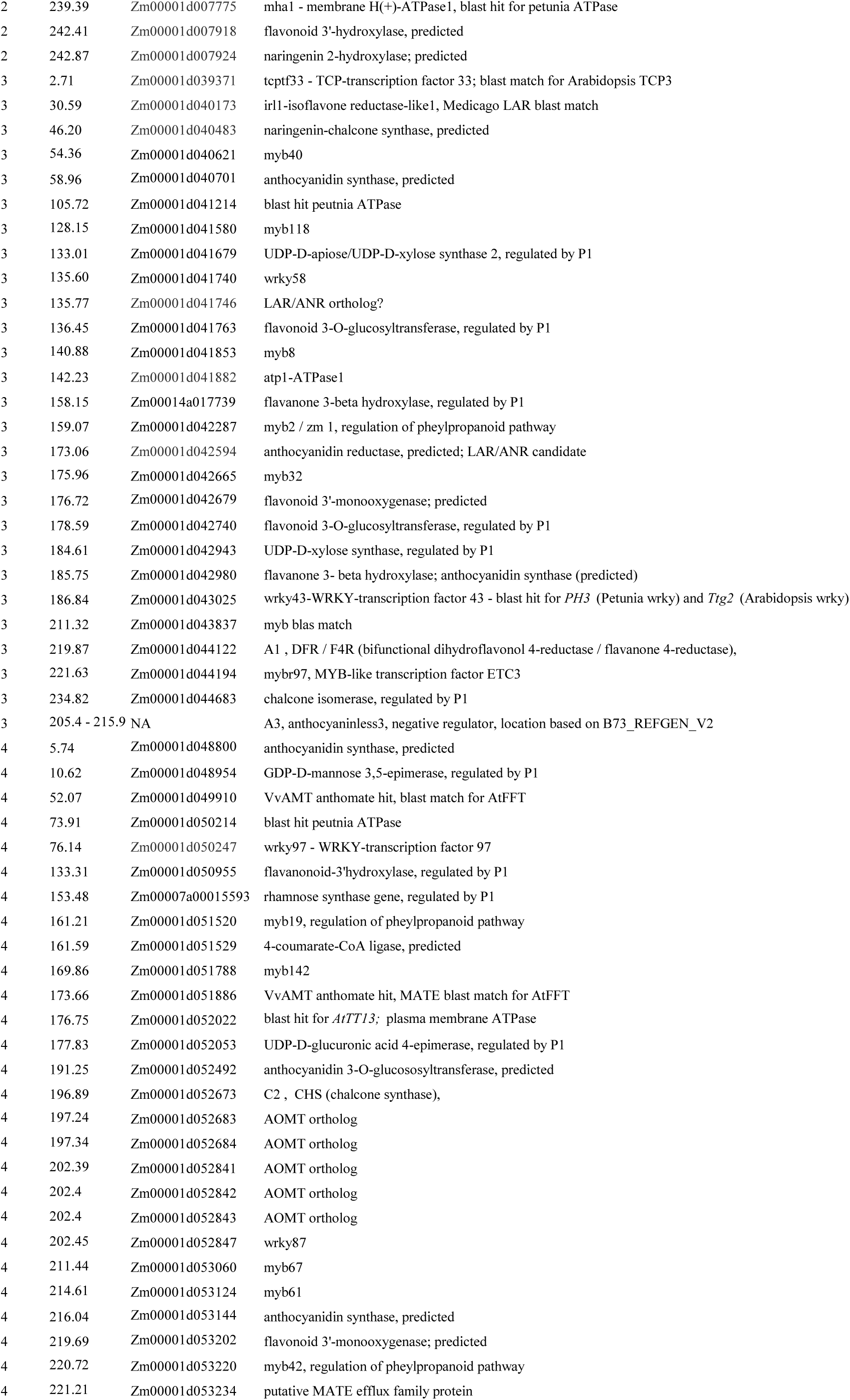
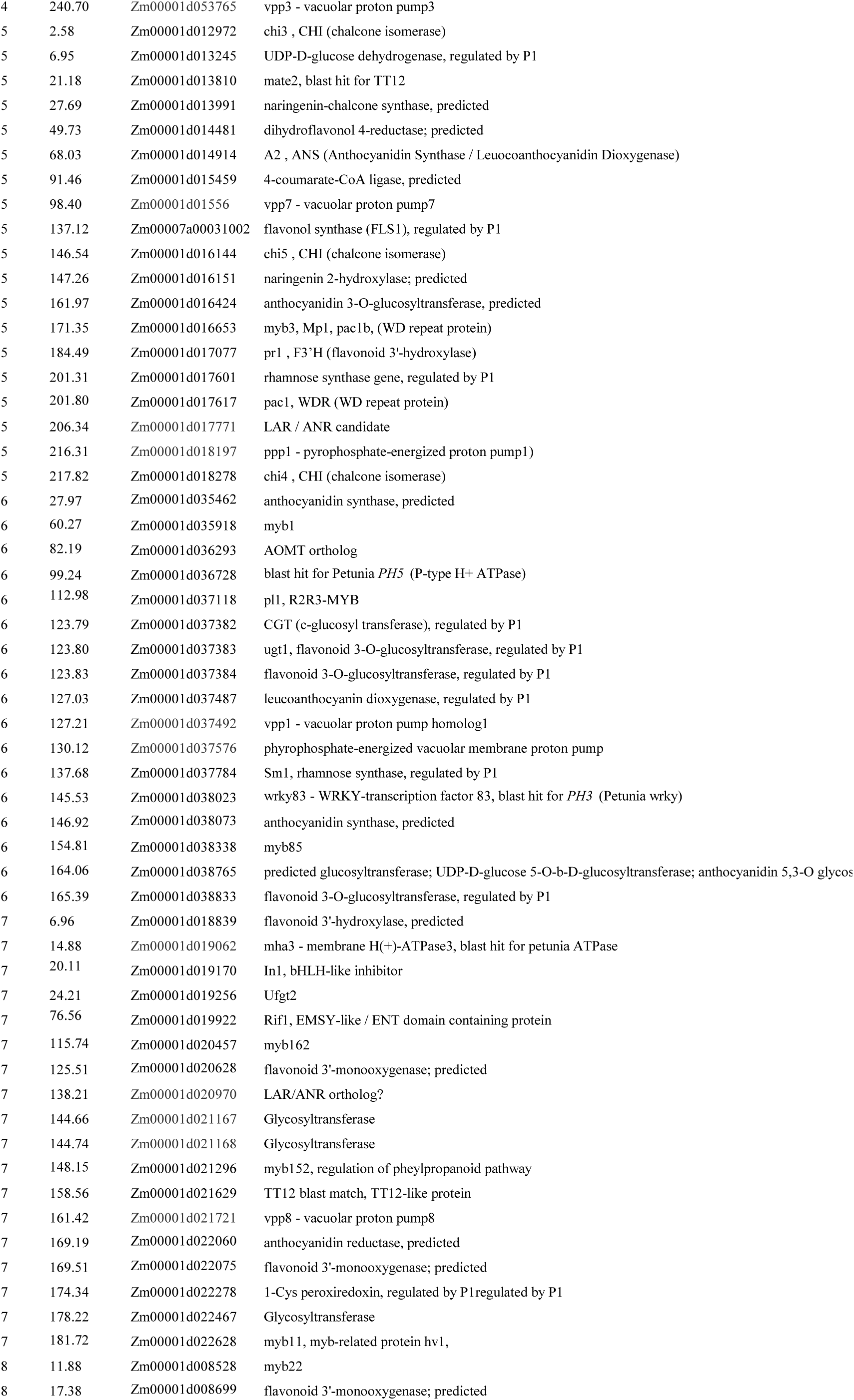
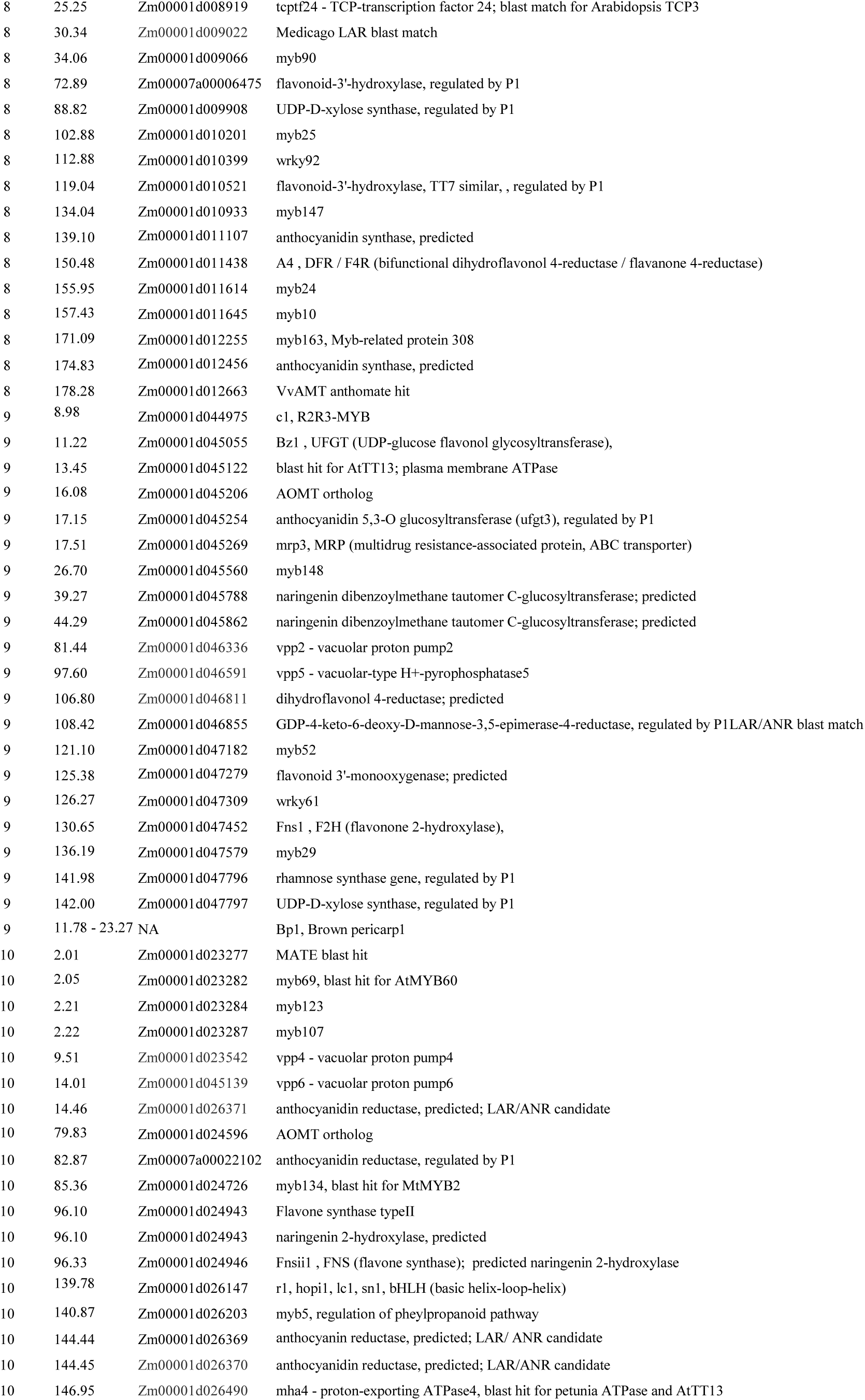
List of potential gene candidates.

**Supplemental Figure 1:**
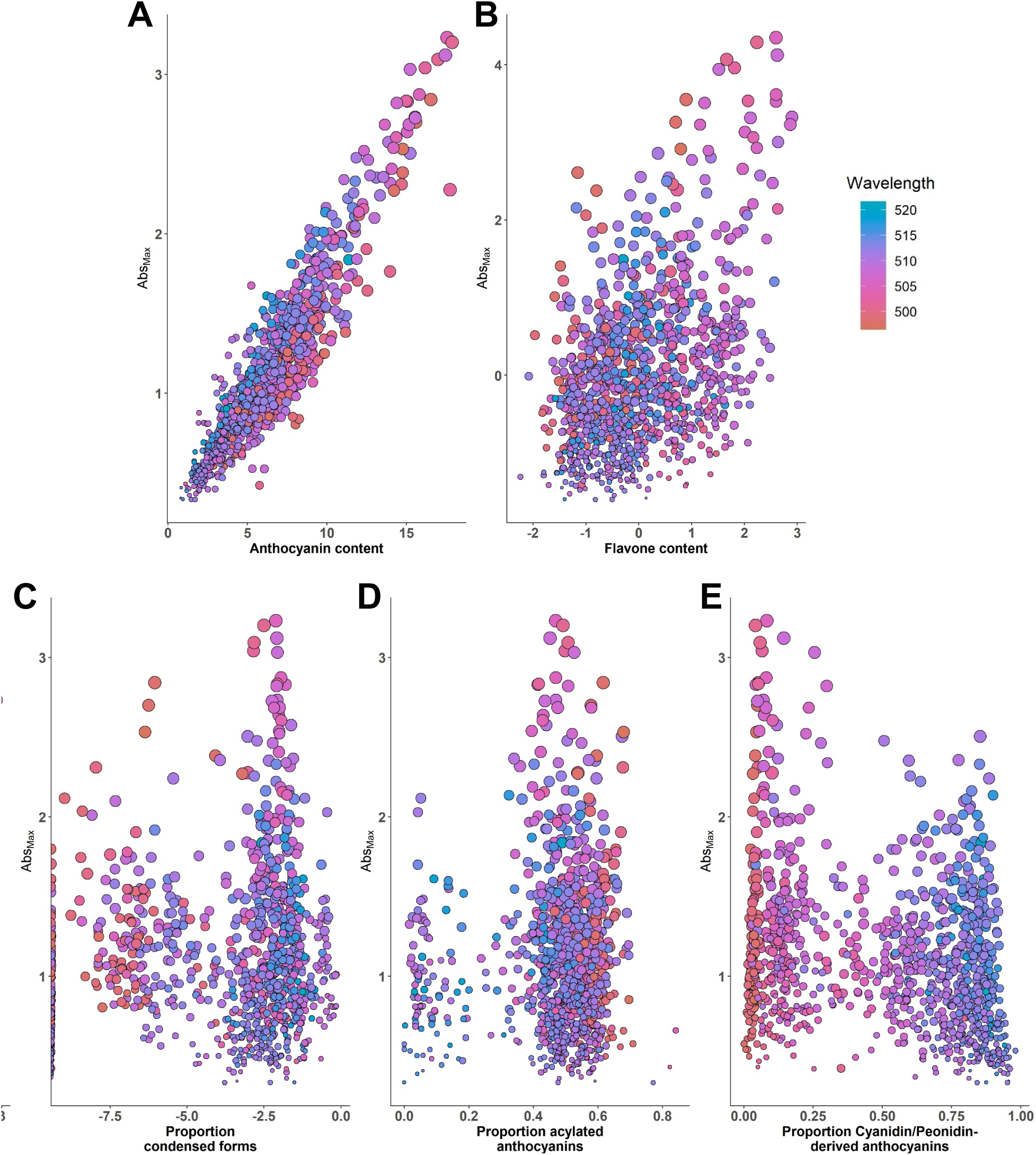
Effects of anthocyanin composition on maximum absorbance (Abs_Max_), a proxy for color intensity. Anthocyanin content was square root transformed from μg/ml, and flavone content (μg/ml) and condensed form proportion were log transformed.

**Supplementary Figure 2:**
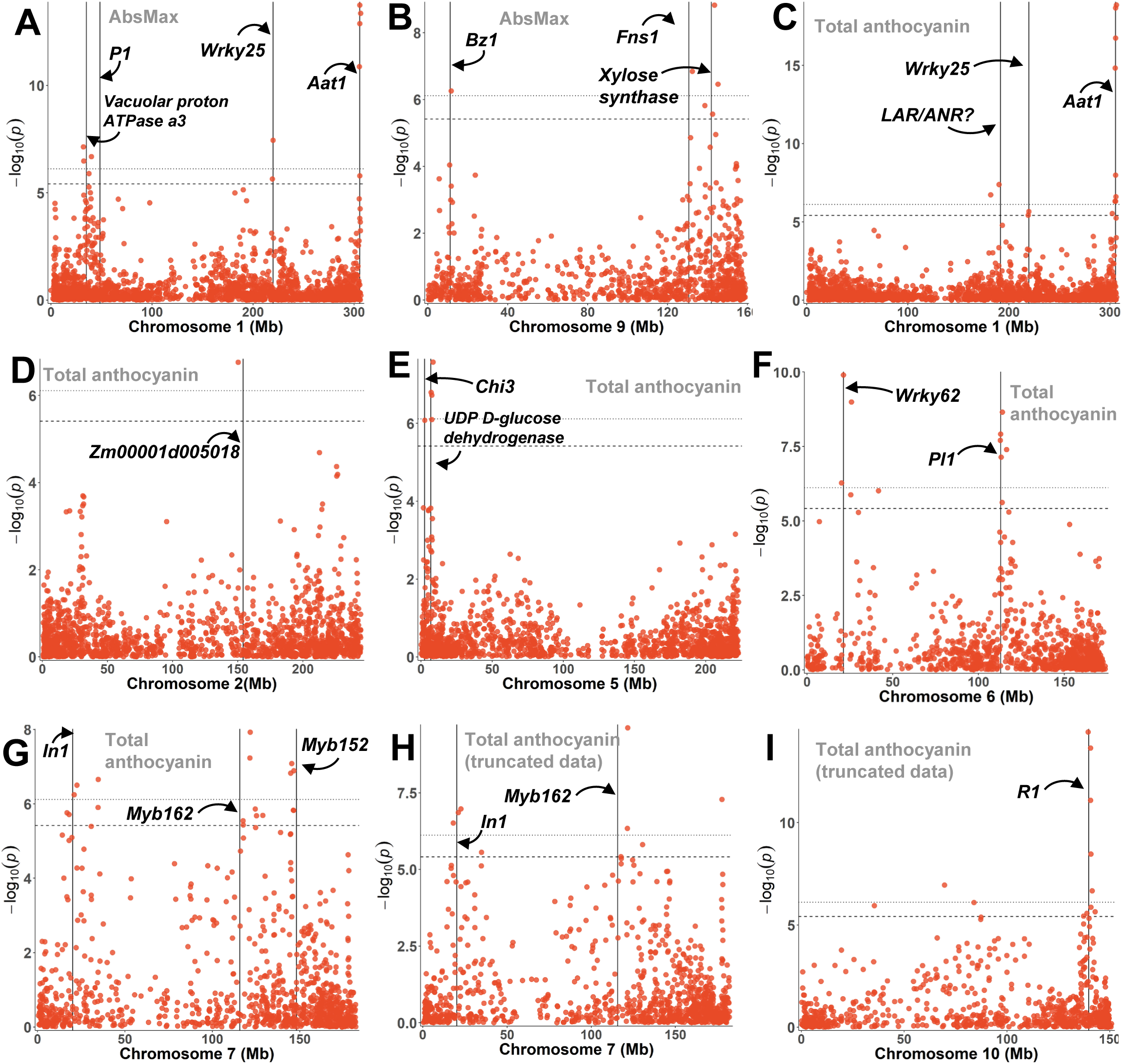
Individual chromosome plots for major signals identified for total absorbance (A, B), anthocyanin content (C-G), and anthocyanin content for the truncated data set (H,I; tails of percent acylation distribution removed). Chromosome numbers are listed in the x-axis label for each plot.

**Supplementary Figure 3:**
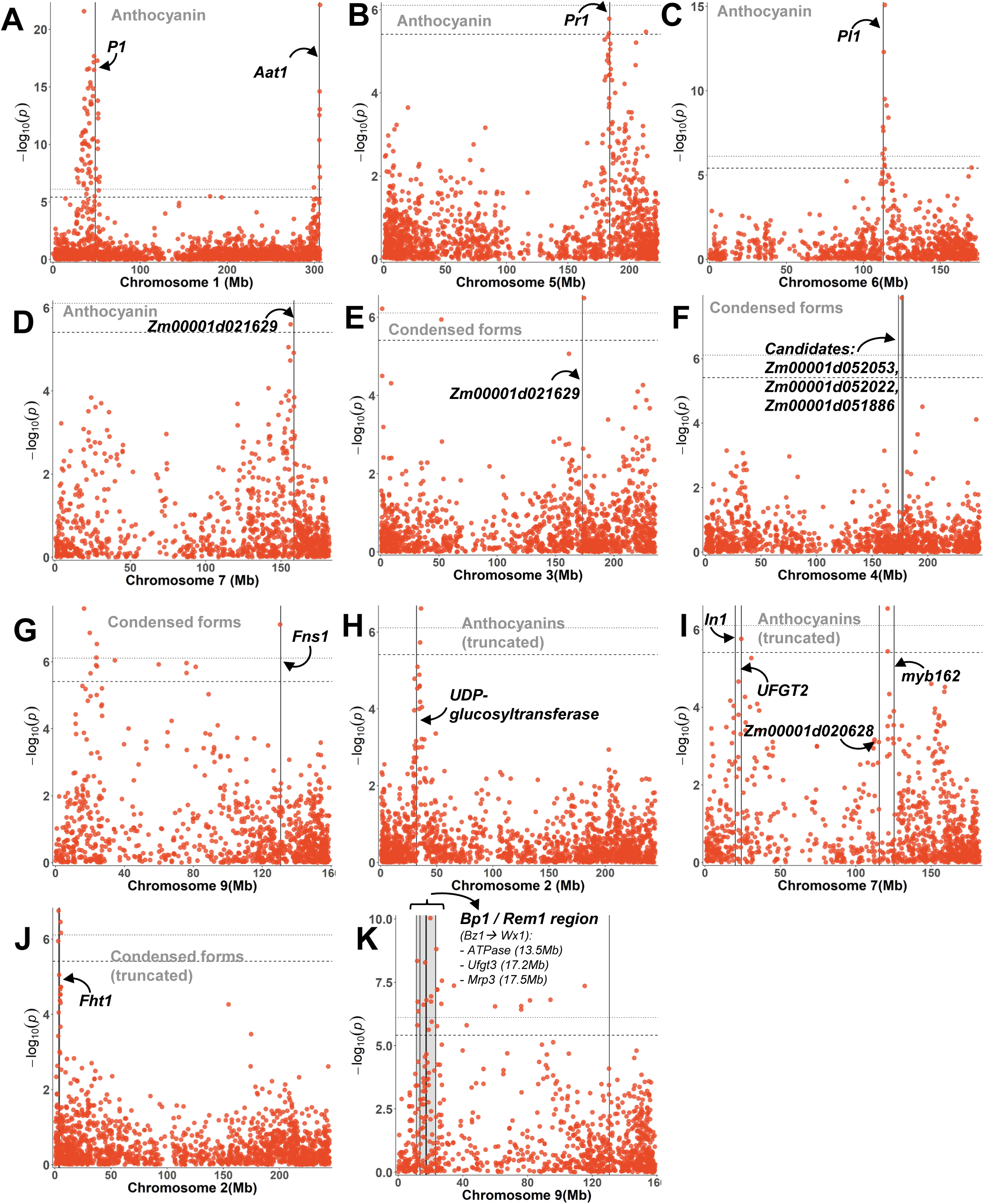
Individual chromosome plots for major signals identified when mapping molar proportions of anthocyanins (A-D) and condensed forms (E-G) in the full data set and for the truncated data set (H-K). Chromosome numbers are listed in the x-axis label for each plot.

**Supplementary Figure 4:**
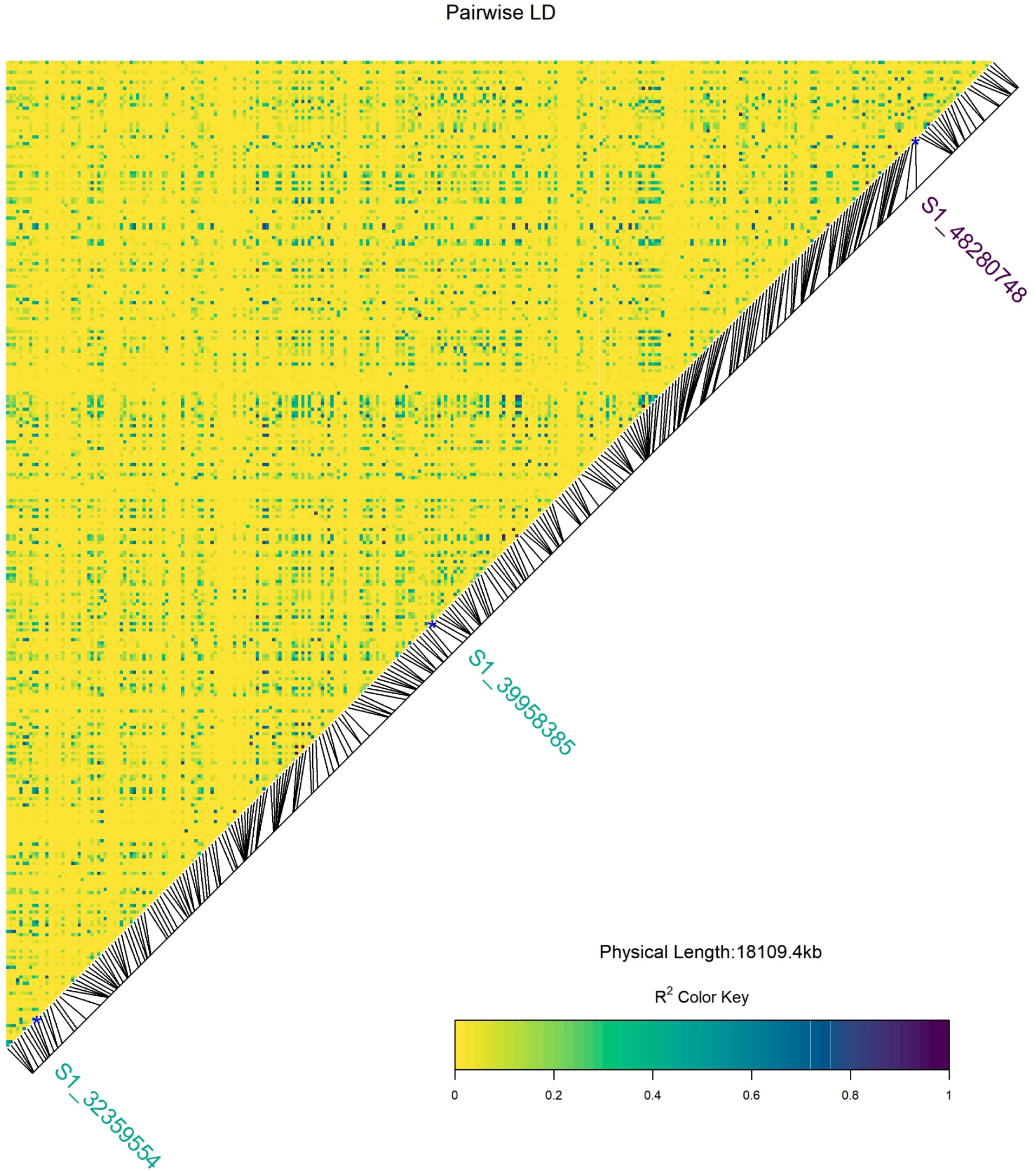
Pairwise LD for chromosome 1 from 33 to 50 Mb. SNPs labeled in green were highly significant SNPs from AR GWAS and the SNP labeled in dark purple is the SNP closest to *P1*.

**Supplementary Figure 5:**
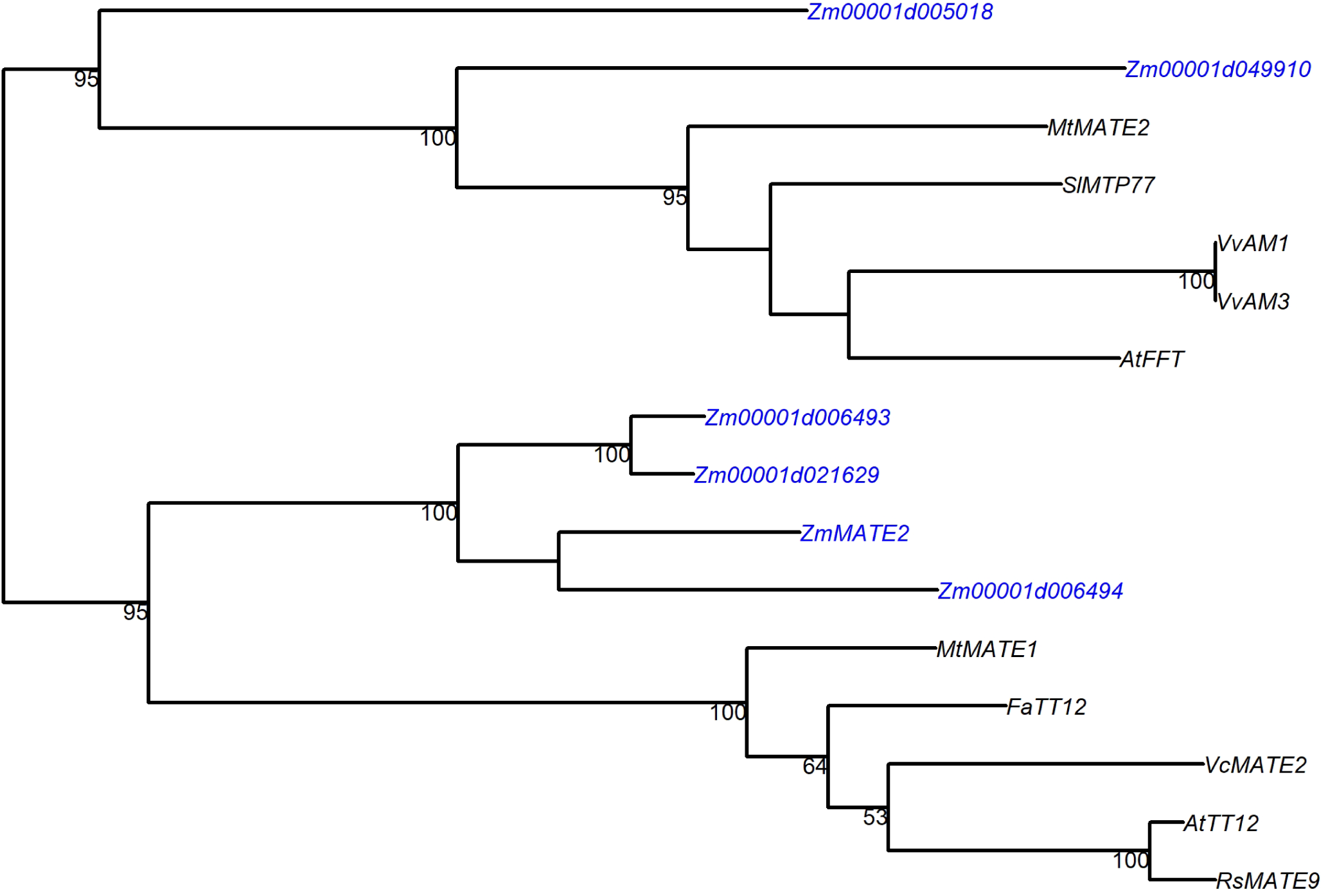
Maximum likelihood phylogeny based on peptide sequences of flavonoid transporting MATEs and candidate MATEs from maize. Branch numbers represent bootstrap values based on 100 replicates.

**Supplementary Figure 6:**
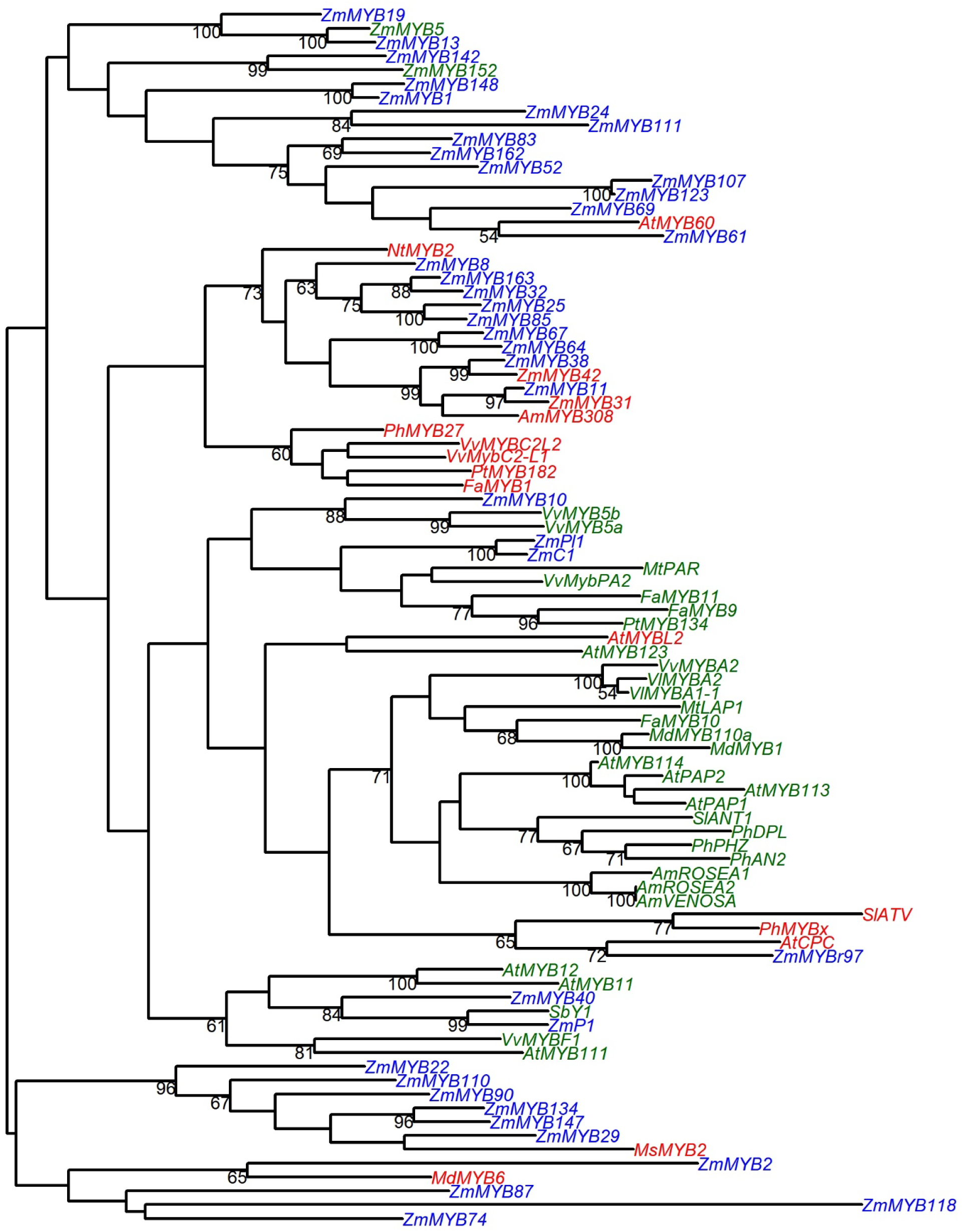
Phylogram of MYBs. Red labels indicate negative regulation while green labels indicate positive regulation of anthocyanin or flavonoid biosynthesis. Maize MYBs with unknown function are colored in blue. Sequences correspond to Supplementary Table 3, and branch numbers represent bootstrap values based on 100 replicates.

